# Food resources drive rodent population demography mediated by seasonality and inter-specific competition

**DOI:** 10.1101/2022.05.07.491005

**Authors:** Ferrari Giulia, Devineau Olivier, Tagliapietra Valentina, Johnsen Kaja, Federico Ossi, Cagnacci Francesca

## Abstract

1. As fast reproducing species, rodents present proximate numerical responses to resource availability that have been assessed by experimental manipulation of food, with contrasting results. Other intrinsic and extrinsic factors, such as climate severity, species life cycles, and sympatry of potential competitors in the community, may interplay to modulate such responses, but their effects have rarely been evaluated *ensemble*.
2. We applied a niche-based approach to experimentally determine the effect of bottom-up (food availability) and top-down (climate severity) extrinsic factors, as well as intrinsic seasonal cycles, on rodent demography, also in presence of sympatry between species in the community.
3. To this end, we live-trapped rodents at two latitudinal extremes of the boreal-temperate gradient (Italian Alps and Norway) deploying control/treatment designs of food manipulation. We applied a multistate open robust design model to estimate population patterns and survival rate.
4. Yellow-necked and wood mouse (*Apodemus* spp.) were sympatric with bank vole in Italy, while the latter was the only species trapped in Norway. At northern latitudes, where harsher climatic conditions occurred, vole survival was principally regulated by intrinsic seasonal cycles, with a positive effect of food also on population abundance. At southern latitudes, mice and voles exhibited asynchronous population patterns across seasons, with survival depending from seasonal cycles. When concentrated *ad libitum* food was experimentally provided, though, population size and survival of voles strongly decreased, while mice abundance benefited from food supplementation.
5. Our results evidence that rodent demography is regulated by a combination of top-down (climate severity) and bottom-up (food availability) extrinsic factors, together with intrinsic seasonal ones. Moreover, we showed that the seasonal niche partitioning of mice and voles could be disrupted by availability of abundant resources that favour the demography of the more opportunistic *Apodemus* spp. at the expense of *Myodes glareolus*, suggesting competitive mechanisms. We conclude putting our results in the context of climate change, where shifts in vegetation productivity may affect the diversity of the rodent community via demographic effects.

## 1 Introduction

Mammalian populations can be placed along a fast-slow continuum according to their life-history traits (Oli, 2004), metabolism (Lovegrove, 2003) and fertility rate (Oli & Dobson, 2003). Extrinsic factors (bottom-up e.g. food availability, top-down e.g. climate severity or predation), together with intrinsic mechanisms (seasonal cycles, density-dependence), control the dynamics of both large, ‘slow-living’, and small, ‘fast-living’ mammals (Odden et al., 2014; Wolff, 1997). However, large mammals have lower basal metabolic rate being related with body mass (Schmidt-Nielsen & Knut, 1984; C. R. White & Kearney, 2013), so that extrinsic perturbations tend to have a delayed effect on individual fitness (e.g. carry-over effect, Harrison et al. 2011; metabolic plasticity, Norin & Metcalfe 2019), hence demography (elasticity patterns, Heppell et al., 2000). Conversely, small mammals show an r-selection strategy with a rapid pace in life-history events (e.g., earlier maturity, higher reproductive rate, shorter generation times and higher basal metabolic rate) (Boyce, 1984), therefore experiencing proximate functional and numerical responses to climatic and environmental changes, such as masting (Zwolak, Bogdziewicz, & Rychlik, 2016), and habitat fragmentation (Bogdziewicz & Zwolak, 2014). For these reasons and due to their wide distribution across diverse habitats and numerous trophic niches, small mammals are considered informative models to disentangle environmental effects on individual fitness, demography and, ultimately, population and community dynamics.

The range of abiotic conditions defining the distribution of small mammal species in absence of other species define their *fundamental niche* (*sensu* Hutchinson, 1957), while in presence of interspecific interactions single populations occur in more restricted ‘regions’ of such environmental space (*realised niche*; see e.g. Peters et al., 2013). Within these regions, population growth rate, expressed by its demographic components such as survival, somatic growth and fecundity (*demographic niche*, Maguire, 1973) depends on the acquisition of food resources, in presence of competitors, predation and diseases (Chase & Leibold, 2003; Sibly & Hone, 2002). In particular, Sinclair and Krebs (2002) suggested that food supply is the primary factor determining rodent population growth rate and only secondarily it is overridden or severely modified by regulatory top-down processes (e.g. predators, intra-specific social interactions and environmental disturbance such as climate severity). This conjecture was then revised by Krebs (2013), proposing instead a multi-factorial explanation of population dynamics changes as an interaction of top-down and bottom-up mechanisms. More recently Flowerdew et al. (2017) supported the earlier model in which food availability, together with density-dependence, play the strongest role in rodent population growth and reproduction, mediated by weather seasonality and only weakly by predation.

A comprehensive analysis of the factors that regulate population dynamics in small rodents is therefore extremely complex because both reproduction and survival are condition-dependent and therefore regulated directly by intrinsic and extrinsic factors. Specifically, intrinsic factors (i.e. density-dependence, behaviour, physiology and life-history traits) cause variations in survival (Johnsen et al., 2018) and reproduction (Aars & Ims, 2002) year-round. Their effect may then interact with environmental extrinsic conditions, such as climatic constraints (Hoset et al., 2009) and food quality and quantity (Boutin, 1990), to shape small rodents demographic patterns.

Several studies evaluating the proximate importance of food in regulating seasonal demography and multiannual population dynamics used supplemental food provisioning as an experimental approach to control food availability (e. g. Boutin, 1990). These studies found that *ad libitum* supplemental food favored breeding (Krebs & DeLong, 1965), increased reproduction (Desy & Thompson, 1983) and promoted survival (Johnsen et al., 2016) leading to an increase in population density (Taitt & Krebs, 1981; Yoccoz et al., 2001). However, in alternate studies food supplementation decreased survival (Hansen & Batzli, 1978), population density (Krebs & DeLong, 1965), and growth rate (Löfgren et al., 1996).

Thus, previous literature was not conclusive in outlining the relationship between food availability and population parameters and patterns, in small mammals. This relationship might be further complicated by interspecific interactions (e.g. Brown & Munger, 1985). Indeed, when ecologically similar species are sympatric, they can either (i) adjust their foraging tactics, habitat selection or activity patterns (i.e. specialist to generalist and opportunistic continuum) to partition the niche in space or time, and thus facilitate coexistence (Leimgruber et al., 2014; Schoener, 1974); or (ii) get involved in competitive mechanisms, such as exploitative competition and behavioural interference that can affect demography parameters of outcompeting subordinate species (e.g., dominant mice aggressiveness *vs* voles, (Bartolommei et al., 2018; Grüm & Bujalska, 2000), ultimately leading to the formation of new community assemblies (Chesson & Kuang, 2008; Eccard & Ylönen, 2003; Price, 1986).

A potential way to explore such complex relationship is by integrating experimental food supplementation designs within a niche-based community approach, where the effects of top-down extrinsic factors, such as climatic constraints, intrinsic seasonal variations, and food availability on demographic parameters of sympatric species are assessed *ensemble*. In this study, we aimed to disentangle the effect of food availability and climate severity, as well as seasonality on small rodents’ demographic patterns, by contemporarily applying a longitudinal and manipulative experimental approach. Specifically, we manipulated food resources availability (i.e., supplemental feeding in treatment/control designs), while explicitly controlling for seasonal variation, degree of climatic severity and co-occurrence of sympatric species, in wild settings. Small rodents were intensively live-trapped at two latitudinal extremes of the boreal-temperate climatic gradient (South-East Norway and Italian Alps), as proxies for climatic severity. In particular, the Norwegian study area is characterized by a strong seasonality with cold and snowy winters and mild and short summers; conversely, in the Italian study area the seasonal shift is more gradual with short and cool winters and warm and flourishing summers. Our initial hypothesis stated that rodents’ population parameters and size would vary with resource availability, seasonal variation, and climatic severity (H1; Table1). In particular, for each detected small mammal species, we expected a general decrease of survival at northern latitudes where conditions are harsher (P1.1; Table 1) whereas, within each study area, survival would depend on species-specific intrinsic factors, such as seasonal physiological cycles (P1.2a; Table 1), and be relatively independent from supplemental food (P1.2b; Table 1). Further, we predicted population size to be positively affected by resources availability at both latitudes, i.e. to be higher in summer (P1.3a; Table 1), especially where supplemental food was available (P1.3b; Table 1). We further hypothesized that rodent assemblage and inter-specific relationships depend on climatic and environmental conditions (H2, Table 1) that when favorable, as at southern latitudes promote a more diversified rodent assemblage (P2.1; Table 1). We also predicted that competitive mechanisms between dominant and subordinate species of the assemblage (e.g. mice *vs* voles) would emerge especially under favorable conditions for dominant species, such as in summer (P2.2a; Table 1) or in presence of supplemental food (P2.2b; Table 1).

**Table 1:**
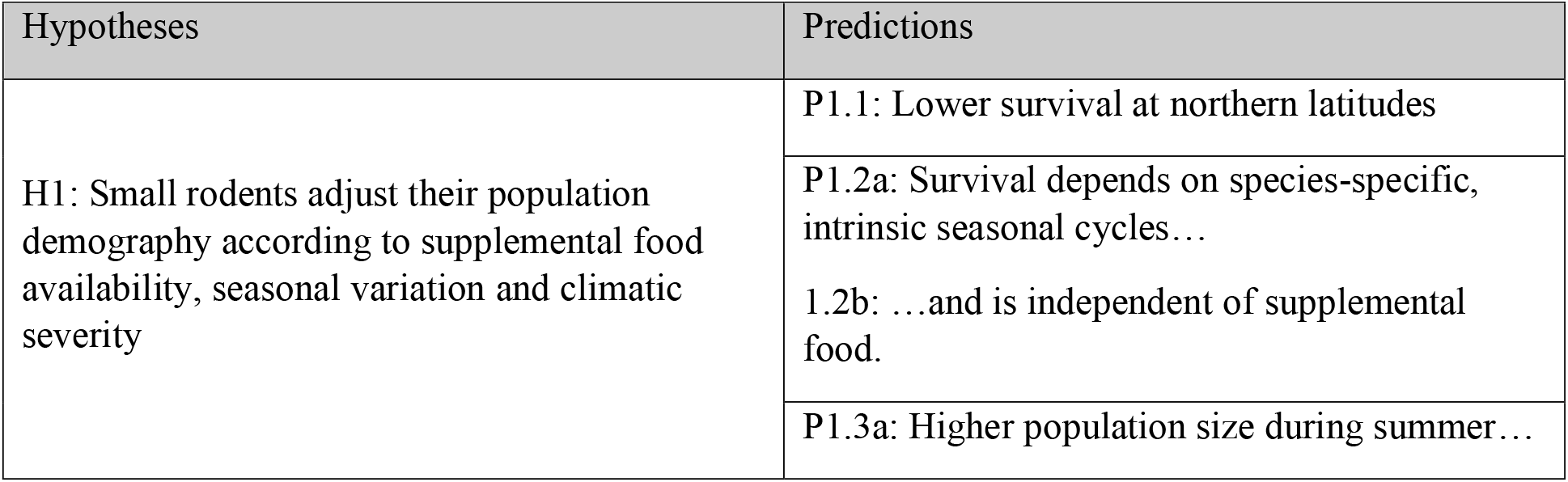
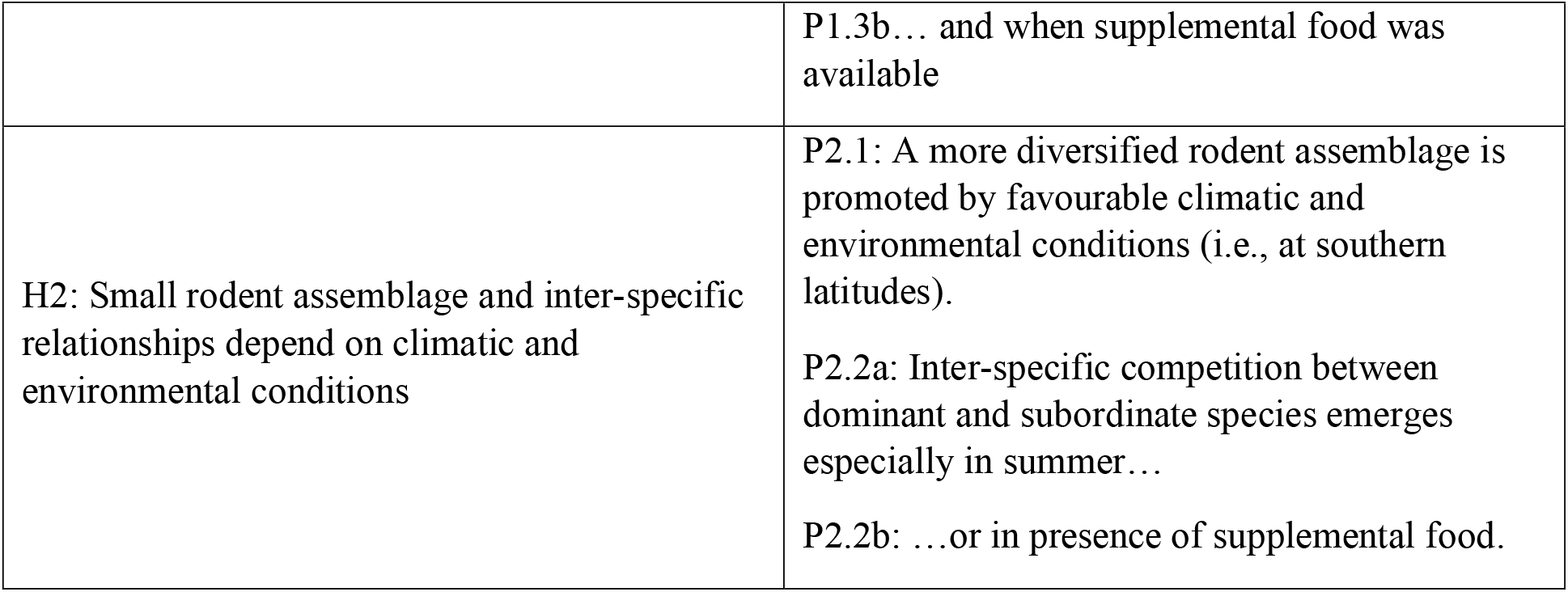
Hypotheses and corresponding predictions.

| Hypotheses | Predictions |
| --- | --- |
| H1: Small rodents adjust their population demography according to supplemental food availability, seasonal variation and climatic severity | P1.1: Lower survival at northern latitudes |
|  | P1.2a: Survival depends on species-specific, intrinsic seasonal cycles... |
|  | 1.2b: ...and is independent of supplemental food. |
|  | P1.3a: Higher population size during summer... |
|  | P1.3b... and when supplemental food was available |
| H2: Small rodent assemblage and inter-specific relationships depend on climatic and environmental conditions | P2.1: A more diversified rodent assemblage is promoted by favourable climatic and environmental conditions (i.e., at southern latitudes).<br>P2.2a: Inter-specific competition between dominant and subordinate species emerges especially in summer...<br>P2.2b: ...or in presence of supplemental food. |

## 2 Materials and Methods

### 2.1 Study areas

We carried out the study in two areas located at the extremes of the boreal-temperate climatic gradient, namely South-East Norway, and North-Eastern Italian Alps (Figure 1).

**Figure 1:**
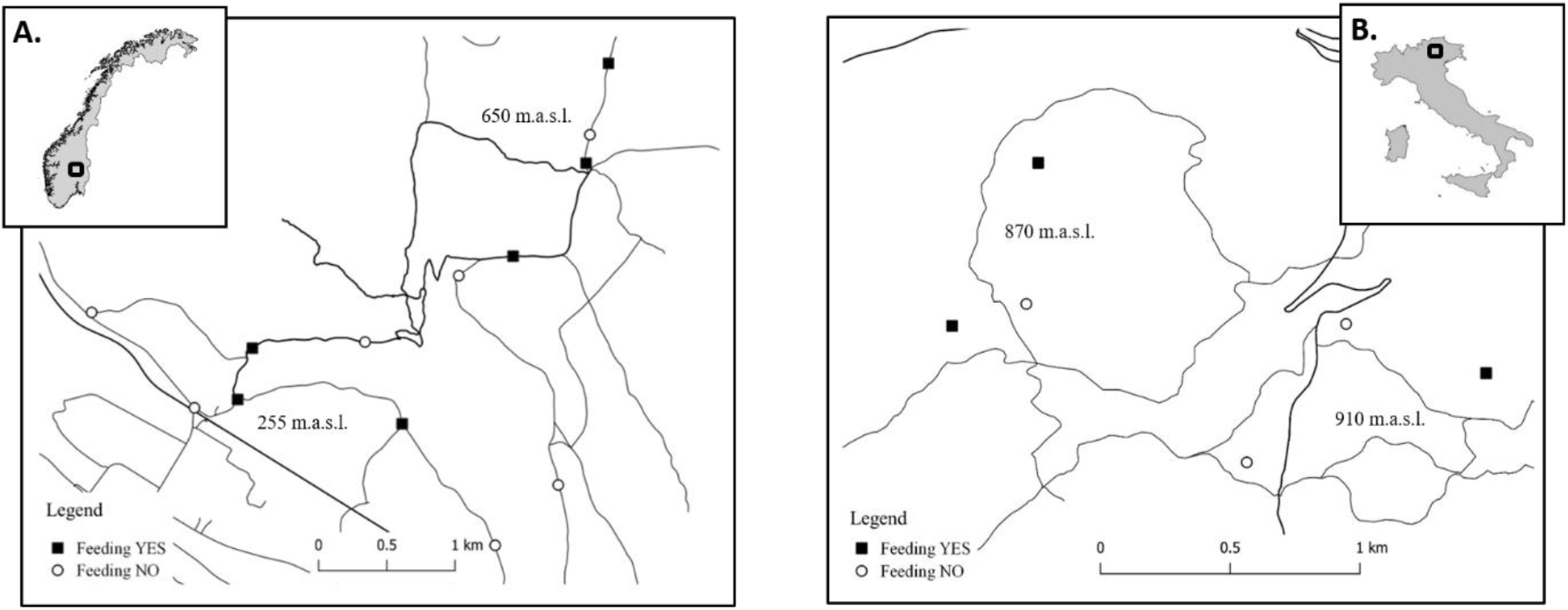
Maps and locations of the study areas. A: Evenstad area, in South-Eastern Norway (61.00N –11.00E), monitoring period: 2013-2015; and B: Cembra area, in North-Eastern Italian Alps (46.13020N – 11.17843E), monitoring period: 2019-2021. In both study areas, rodents were live-trapped at grids with experimental control-treatment settings regarding food resource availability. Thick lines in the bottom valley indicate the main roads and fine lines the forest roads.

In Norway, Evenstad study area (61.00N – 11.00E; Stor Evdal municipality) includes the forested areas covering the hills and the valley bottom of the Glomma river basin (255 – 700 m a.s.l.). It is characterised by boreal climate (*sensu* Köppen-Geiger classification, Kottek et al., 2006), with long, harsh and snowy winters, with a permanent snow layer that prevents the soil from freezing, and short, cool summers. In particular, mean daily temperature of –8.9 ± 3.2 °C in January and of 14.2 ± 2.1 °C in July, and mean monthly precipitation of 35.2 ± 14.7 mm in January and of 84.1 ± 40.4 mm in July were recorded (data of 2000 – 2020 from the Norwegian Meteorological Institute https://seklima.met.no/, Evenstad, Åkrestrømmen, Drevsjø and Gløtvola weather stations at 260 / 670 m a.s.l.). The landscape is dominated by homogenous boreal coniferous forest of *Picea abies* and *Pinus sylvestris*, bilberry (*Vaccinium myrtillus*) in the understory shrub layer, and mosses (e.g., *Pleurozium schreberi*) on the ground layer (Johnsen et al., 2016).

In Italy, Cembra Valley (46.13020N – 11.17843E; Autonomous Province of Trento, Albiano municipality) lays in a moderate topography region (500 to 1000 m a.s.l.) in orographic continuity with the Lagorai massif. It is characterised by warm-temperate class of the alpine climate, *sensu* Köppen-Geiger classification (Rubel et al., 2017), with moderately cold winters, with occasional snow cover and usually frozen ground, and fresh summers. In particular, mean daily temperature of 0.9 ± 1.3 °C in January and of 20.8 ± 1.4 °C in July, and mean monthly precipitation of 39.0 ± 1.3 mm in January and of 112.9 ± 47.8 mm in July were recorded (2000 –2020 data obtained from Meteotrentino https://www.meteotrentino.it, Cembra weather station at 652 m a.s.l.). The area is covered with relatively homogeneous secondary growth mixed forest, dominated by *Pinus sylvestris* with abundant shrub undergrowth, as well as mixed stands of *Fagus sylvatica, Picea abies, Abies alba* and, to a lower extent, *Quercus petraea*, interspersed with peat bogs and small pastures. In 2020, a mast seeding event of both *Fagus sylvatica* and *Picea abies* occurred in the Italian study area (Ferrari G, pers. comm.).

### 2.2 Experimental designs

In this study, we assessed rodent demography (i.e. survival and population size) under contrasting abiotic conditions (climatic severity and seasonal variation) and food availability (H1). In particular for the former, we compared (i) study areas at latitudinal extremes, namely Norway and Italy, as a proxy for climatic severity, and (ii) permissive (summer: April-October) vs limiting (winter: November-March) periods, within each year, to evaluate the impact of seasonal variation on rodents demography. To evaluate the demographic implications of food availability, we undertook experimental manipulations by providing *ad libitum* supplemental food in some trapping occasions and/or sites (‘Feeding yes’, or ‘Treatment’ Figure 1, Figure 2), but not in others (‘Feeding no’, or ‘Control’ Figure 1, Figure 2), in both study areas. Additionally, we further investigated rodent adaptability to divergent environmental conditions by comparing the assemblages at the two latitudes (H2). Where sympatric rodent species occurred, we assessed whether they were competing by comparing their demographic parameters under the aforementioned seasonal and supplemental food availability variations (Figure 2).

**Figure 2:**
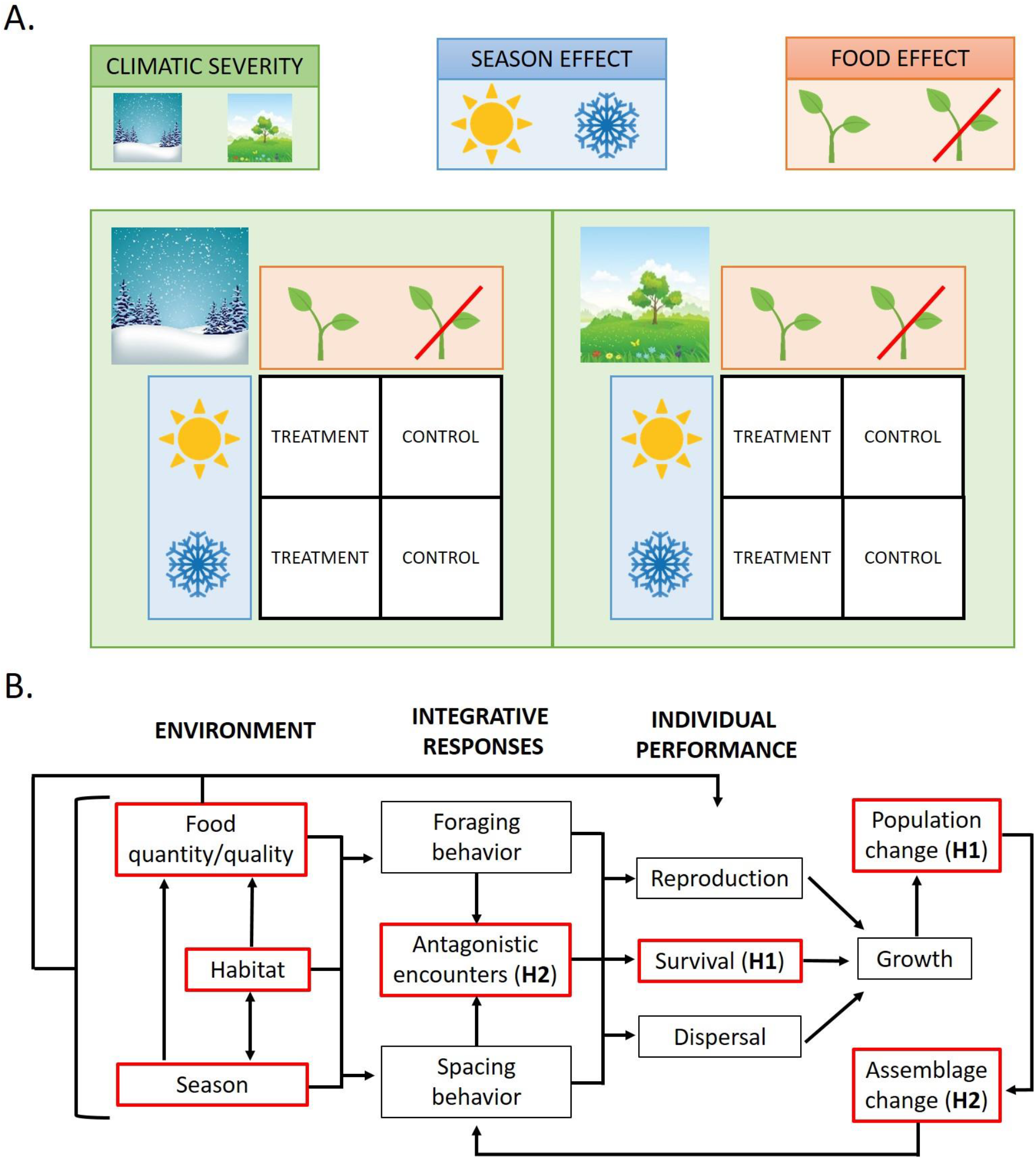
Panel A - In this study, we performed rodent live-trapping at two latitudinal extremes (Norway and Italy) used as a proxy for climatic severity (in green), in a combination of seasonal (in blue) and supplemental food availability (in orange) conditions. Specifically, at both latitudes, we performed experimental control-treatment manipulations in wild settings by providing supplemental food in treatment conditions only, across different seasons. Panel B - These experimental manipulations led to evaluate individual performance and population abundance change in dependence on seasonal variation and food availability, under diverse degree of climatic severity (H1). Additionally, we evaluated how these relations were affected by competition of sympatric species (H2). Scheme modified from (Batzli, 1992); the red boxes indicate the factors explicitly addressed in the study designs.

Specifically, in Norway, we deployed thirteen trapping grids and performed monthly captures between 2013 and 2015 (see Appendix S1, Table S1.1 for a summary of capture sessions). In six of those grids, supplemental food (a mixture of oat and sunflowers) was provided *ad libitum* next to each trap from December 2013 to June 2014, in November 2014, and in May 2015 (state of the sites: ‘Feeding yes’). In these six grids from June to August 2015, and in the remaining seven grids from December 2013 to August 2015, no food was provided (state of the sites: ‘Feeding no’).

In Italy, we deployed six trapping grids and performed monthly captures during winter (from November to March), and bi-monthly captures during spring and summer (i.e., in April, June and August) between February 2019 and April 2021 (see Appendix S1, Table S1.2). Three of these grids were always provided with *ad libitum* corn supply dispensed through ungulate supplemental feeding stations located at the centre of each grid (sites: ‘Feeding yes’), while the other three were located at least 500 m from any feeding site (sites: ‘Feeding no’).

### 2.3 Live-trapping

In Norway, the trapping grids were cross-shaped with 16 trap stations 15 m and 7.5 m apart for the external and internal traps in the grid, respectively (total grid area: 3600 m^2^; Appendix S2). Captures lasted 4 days/3 nights per month, and traps were checked every 12 hours (for additional information on the protocol in Norway, see Johnsen et al., 2017).

In Italy, animals were caught in square-grids of 8×8 traps placed 10 m apart (total grid area: 4900 m^2^; Appendix S2). Captures lasted 4 days/3 nights per month (winter) or bi-month (summer), and traps were checked every 24 hours.

In both the Italian and Norwegian study areas, the traps were left open during non-trapping periods, to habituate animals to their presence. Each trap consisted of a standard Ugglan Multiple Live Trap (model 2, Granhab, Sweden), filled with hay during the cold season, and baited with carrot slices and seeds (oat in Norway, sunflower in Italy). Standard capture-mark-recapture techniques (CMR) were adopted (Amstrup et al., 2005; Pollock et al., 1990). Each animal was individually marked with a Passive Integrated Transponder (PIT) tag (Trovan® Ltd., UK), and standard information (date, time, trap number, grid number, id of the animal) and life-history traits (species, sex, body mass, age, breeding status) were recorded for each capture. In Italy only, blood (from the retro orbital sinus), tissue samples (tail tip and ear biopsy) and fecal pellets were collected, and animals were also screened for the presence of parasites.

We discarded data from 22 animals in total as they were occasional species (shrews, dormice), or individuals that lost their PIT tag (but were identified by the signs of biopsy). Individuals of yellow-necked mouse (*Apodemus flavicollis*) and wood mouse (*Apodemus sylvaticus*) were treated *ensemble* as *Apodemus* spp. as the former was largely prevalent (see below).

All animal handling procedures and ethical issues were approved by the relevant regional or local Wildlife Management Committees.

### 2.4 Multi-State Open Robust Design models

We analysed capture data separately for Norway and Italy to account for differences in the experimental design (see 2.2 *Experimental designs*).

Our field protocol defined by daily secondary occasions within monthly primary ones corresponds to the so-called robust design framework (Pollock, 1982). Therefore, we used a multistate open robust design (MSORD) approach to estimate demographic parameters (Cooch & White, 2019; G. C. White et al., 2006). This model provides five parameters (Boys et al., 2019; Cooch & White, 2019; Kendall & Nichols, 2002): (i) true survival (S_t_), which expresses the probability of surviving from release occasion t to subsequent primary occasion t + 1 occupying a state s; (ii) transition probability (*ψ*_s_), which represents the probability of animals transitioning to different state s s between primary occasions t and t + 1; (iii) entry or arrival probability (*pent*_*j*_), which expresses the probability of an animal arriving and being captured in the study area in secondary occasion j; (iv) apparent survival rate (*φ*_*j*_), which is the probability of an individual to survive and persist in a capture site at secondary occasion j, given it was present in j – 1; (v) capture probability (*p*_*j*_), the probability of an animal being detected at occasion j, given it was present. The transition probability (*ψ*) was intended differently for Norway and Italy. For the former, it consisted in the probability of transition between state s of food availability at trapping sites, expressed by the covariate ‘feeding state’, as feeding was treated as a spatio-temporal covariate (i.e., availability of food in certain primary occasions in certain sites; see also 2.2 *Experimental designs*); for the latter, it regarded the probability of transition between state of individual observability, i.e. animals being present in the study area and available for detection (observable state) or otherwise temporary emigrant outside the study area (unobservable state), and it was set as constant. The availability of supplemental food for Italy was expressed instead by a spatial, temporally fixed covariate, ‘feeding site’, corresponding to the trapping sites where food was always provided *ad libitum*, or the alternate sites with no food, that as such did not affect *ψ*. The MSORD model also provides two derived (i.e. not included in the likelihood formulation) parameters: (vi) population size (*N*_*t*_), i.e. the number of individuals occupying the study area in state s in that primary occasion t; and (vii) residence time (*R*_*t*_), which expresses the average number of secondary occasions that an individual spent in the study area for that state s in that primary occasion t (Boys et al., 2019; Chabanne at al., 2017).

We modelled demographic parameters in dependence on temporal, spatial, state, and individual covariates that were biologically meaningful to address our working hypotheses. Based on exploratory analyses, we decided not to consider sex in the CMR models (see Appendix S3 for details). We modelled true (*S*) and apparent survival (φ) as varying in dependence on primary occasions (‘session’, only for φ) and successive seasonal periods (‘season’) to detect the temporal pattern (H1), supplemental food availability (‘feeding state’ in Norway and ‘feeding site’ in Italy) to identify the food effect (H1), and species (only in Italy) to evaluate interspecific competition (H2). We considered the probabilities of capture (*p*) and arrival (*pent*) to be dependent on temporal variations (primary occasions i.e. ‘session’ for *p*, secondary occasions i.e. ‘time’ for *pent*), seasonal periods (‘season’), supplemental food availability (‘feeding state’-Norway and ‘feeding site’-Italy) and species (only in Italy). We modelled *ψ* in dependence on transition of feeding state for Norway, and kept it constant for Italy (expressing the probability of transition of animal observability in the trapping grid; see above) (Table 1; see also Appendix S4 for description of covariates and models building).

We fitted the models with the RMark package (Laake, 2013) in the R program (R Core Team 2021) and used the AICc (corrected Akaike’s Information Criterion) for model selection (Burnham & Anderson, 2002) with a threshold of ΔAICc ≤ 4 (Appendix S5). Two models for Norway, and three models for Italy were equivalent in terms of AICc; among these, we chose for prediction of parameters the models which retained the covariates that allowed us to better test our hypotheses (Appendix S5).

## 3 Results

### 3.1 Assemblage Description

In Norway, 917 bank voles (*Myodes glareolus*) were captured accounting for 3976 total capture events. Individuals were captured more when supplemental feeding was available (n = 575 for 2880 night traps) than when it was not (n = 455 for 10504 night traps). The sex ratio was similar with and without supplemental food availability (Table S3.1, Appendix S3).

In Italy, 507 animals were captured for three species (P2.1 *supported*): 398 (78.5%) *Apodemus* spp. (specifically, 390 yellow-necked mice *Apodemus flavicollis* and 8 wood mice *Apodemus sylvaticus*, as later determined through genetics) and 109 (21.5%) bank voles (*M. glareolus*), accounting for 1379 capture events in total (926 and 453, respectively). We captured more mice where supplemental food was available (n = 234 for 21888 night traps) than in grids where it was not (n = 164 for 21888 night traps), and the opposite for voles (29 and 80, respectively). Sex ratio was similar with and without supplemental food (see Table S3.1, Appendix S3).

### 3.2 Population Demography

#### Probability of survival

Generally, *S* in *M glareolus* was marginally lower in Norway during the winter season compared to Italy (t-test; p-value = 0.051), instead during summer *S* was not statistically different in the two study areas (P1.1, *partially supported*).

For Norway, *S* depended on the additive effect between availability of supplemental food and season (Table S5.1). In particular, *S* was higher during winter with respect to the following summer (for those years where these seasons were comparable) (P1.2a *supported*), and in presence of supplemental food, both in winter and summer (Table S6.1; Figure 3.1; for *p, φ* and *pent*, see Appendix S7) (P1.2b *not supported*).

**Figure 3:**
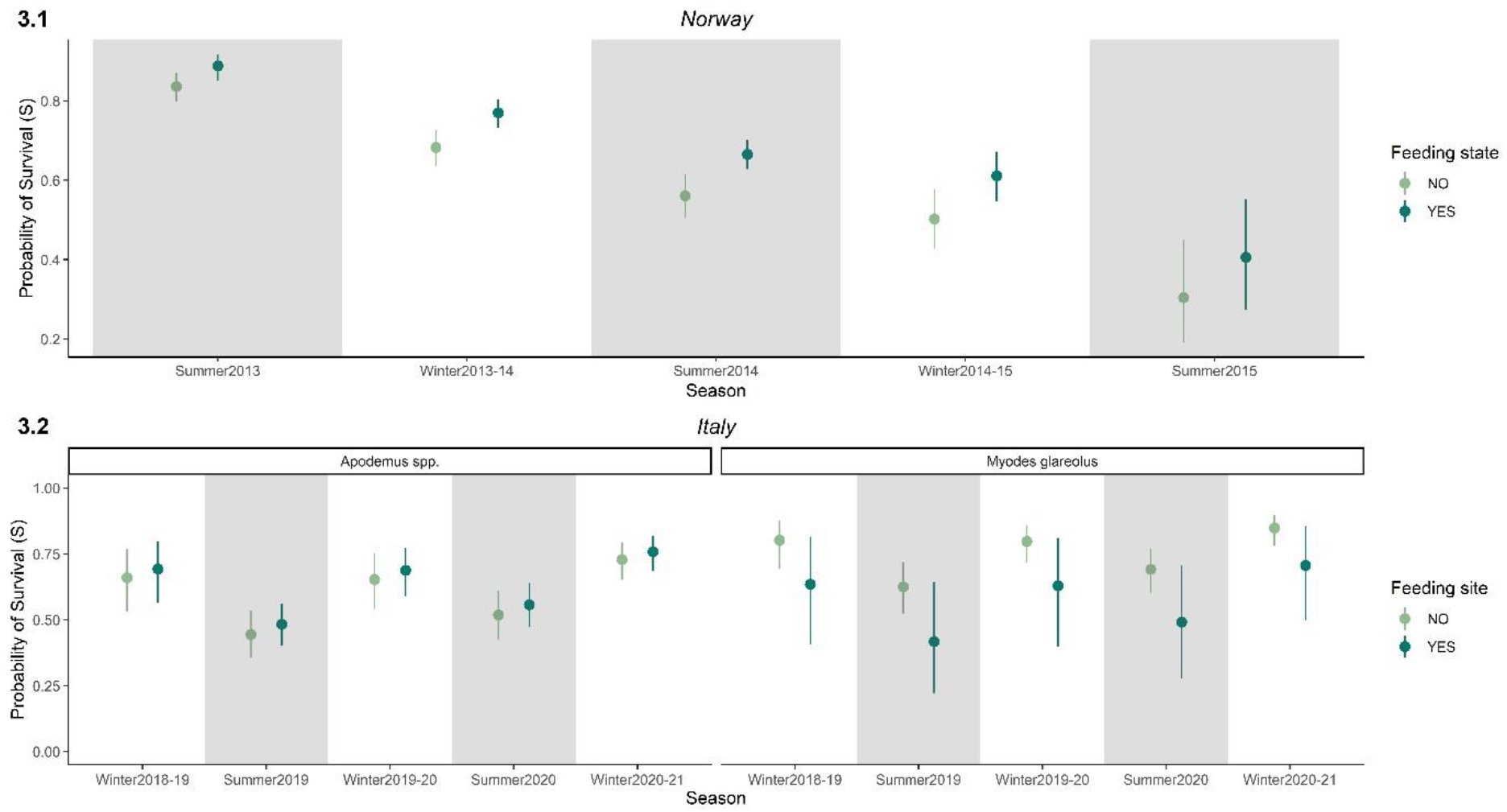
Estimates of survival (S) of rodents in Norway (**Figure 3.1**) and Italy (**Figure 3.2**) by species and seasonal variation, with treatment supplemental food (supplemental food available; dark green) and control (no supplemental food available; light green) conditions ('Feeding state' for Norway and 'Feeding site' for Italy). Grey shaded bars show the summer periods, while white bars show the winter periods. The comparison between successive winter (overwintering animals) and summer (reproductive season) survival estimates is not possible for summer 2013 in Norway and winter 2020-21 in Italy.

In Italy *S* depended on the additive effect of season with the interaction between species and feeding site (Table S5.2). In particular, both *M. glareolus* and *Apodemus* spp. showed a similar seasonal pattern, with *S* increasing in winter and declining in summer (P1.2a *supported*). In absence of supplemental food, *S* was higher in *M. glareolus* than in *Apodemus* spp.; however, where supplemental food was provided, *S* slightly increased in *Apodemus* spp. but strongly decreased in *M. glareolus* (Table S6.3; Figure 3.2; for *p, φ* and *pent*, see Appendix S7) (P1.2b *partially supported* and P2.2b *supported*).

#### Population size

In Norway (Table S6.2; Figure 4.1), when supplemental food was not provided, *M. glareolus* population size was slightly higher in summer than in winter (P1.3a *supported*), although it exhibited a general decreasing trend across years. In the months with availability of supplemental food, the population size was significantly higher (ANOVA, p-value = 0.00013; P1.3b *supported*), but the temporal irregularity of sampling and therefore estimates (Appendix S1, Appendix S6) does not allow to infer any seasonal pattern.

**Figure 4:**
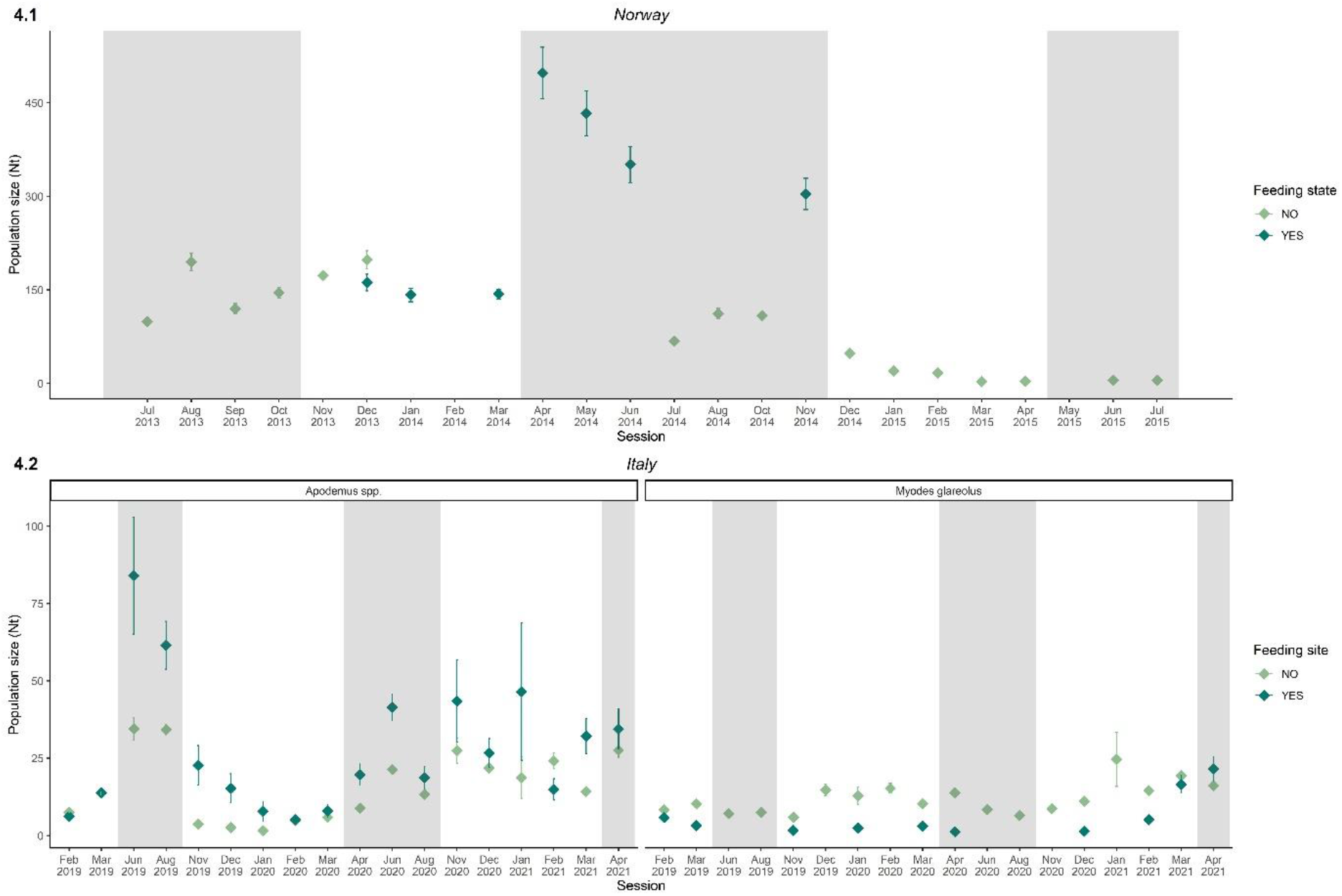
Population size (Nt) of rodents in Norway, 2013-2015 (**Figure 4.1**) and in Italy, 2019-2021 (**Figure 4.2**) with treatment supplemental food (supplemental food available; dark green) and control (no supplemental food available; light green) conditions ('Feeding state' for Norway and 'Feeding site' for Italy). Grey shaded bars show the summer periods, while white bars show the winter periods. The gaps in the series (*M. glareolus* in Norway: Feb 2014 and May 2015; *M. glareolus* in Italy: 8 cases) are due to lack of estimates from the model in those instances.

In Italy (Table S6.4; Figure 4.2), when supplemental food was not provided, population size trends of *Apodemus* spp. and *M. glareolus* were asynchronous, reaching population size peaks in different periods. In particular, *Apodemus* spp. showed a summer peak followed by a sharp decrease through winter 2019-20, while in 2020 the population continued to increase after the summer season (P1.3a *partially supported*). On the contrary, *M. glareolus* increased during winter and decreased in summer (P1.3a *not supported*, P2.2a *supported*). With supplemental food, *Apodemus* spp. exhibited the same annual pattern, but with a higher population size (ANOVA, p-value = 0.03) (P1.3b *supported*). Noteworthy, *M. glareolus* was almost absent where food was provided, with the exception of a slight recovery only during the last winter sessions in 2020-21 (ANOVA, p-value = 0.01) (P1.3b *not supported* and P2.2b *supported*).

## 4 Discussion

In our work, we found evidence that bottom-up extrinsic processes, i.e. food availability, crucially affect woodland rodent demography, and that this effect is mediated by top-down extrinsic drivers, such as climate severity, as well as by intrinsic seasonal cycles and competition between sympatric species. Survival was principally governed by intrinsic seasonal cycles and was affected by food availability only when unfavourable extrinsic conditions occurred (i.e., harsh climate in Norway; exploitative competition in Italy, see below). Conversely, population size was generally enhanced by resources (i.e., with supplemental food in Norway; during masting in Italy), unless *ad libitum* concentrated food availability triggered interactions between sympatric species. In this case, the dominant, opportunistic species (i.e., *Apodemus* spp.) relatively increased its population size both in summer and especially in winter with respect to control conditions, suppressing the temporal niche partitioning with the subordinate generalist species (i.e., *M. glareolus)*. As this resulted in the depression of both population size and survival of the latter, we can infer the emergence of both direct (behavioural interference, i.e. decreased population size) and indirect (exploitative, i.e. depressed survival) competitive mechanisms. Innumerable studies have contributed to describe and explain rodent dynamics by investigating the underpinning demographic processes (Andreassen et al., 2021; Cornulier et al., 2013; Oli, 1999). The niche-based, experimental and longitudinal designs of our work allowed to uncover the interplay between intrinsic and extrinsic drivers of annual population patterns.

Both for bank voles and mice, well-known patterns of seasonal demography for rodents living in temperate and polar climate, i.e. an alternation between breeding (late spring to early autumn) and non-breeding (late autumn to early spring) seasons (Eccard & Herde, 2013) were mainly confirmed. This suggests that intrinsic mechanisms such as physiological cycles represented the leading force in driving survival, which was higher in winter than in summer (P1.2a), and population size that conversely was higher in summer than in winter (P1.3a). Specifically, in late spring, the decreased costs of thermoregulation due to warmer temperatures (Merritt, 2010), together with increased resource availability (Pearse et al., 2016) stimulate reproduction (Steinlechner & Puchalski, 2003), which continues until early autumn producing several generations of new-borns. During the breeding season, the survival is lowered by a combination of (i) high emigration rate of overwintered individuals (i.e. those that successfully survived throughout the previous winter due to their ability to storing food and avoiding predation (Olenev & Grigorkina, 2014) and (ii) deaths of early-born and immature animals (Flowerdew et al., 1985) that tend to increase their competitiveness for food and mates at the cost of chances of survival (Eccard & Herde, 2013). During the non-breeding season, the survival of late-born animals (i.e., overwintering individuals) is higher, guarantying the bulk for the following year spring population.

Our integrated niche-based approach allowed to experimentally evaluate how extrinsic environmental factors, especially trophic resources, modulate these relatively conserved demographic patterns. Specifically, the latitudinal comparison highlighted that *M. glareolus* survival was slightly lower in Norway than in Italy, but only in winter, likely due to prolonged harsh seasons, scarcity of food resources, and delayed phenology (P1.1; Haapakoski & Ylönen, 2013; Korslund & Steen, 2006). Consistently, the survival in *M. glareolus* increased following the manipulation of food availability in Norway (on the contrary of what expected; P1.2b), providing a further evidence of the importance of tropich resources in modulating survival at northern latitudes (Johnsen et al., 2016; Rémy et al., 2013). Indeed, at those latitudes harsh conditions increase the energetic costs for thermoregulation and reproduction (Ylönen & Eccard, 2004). On the opposite, short winters, milder temperatures and more predictable precipitations might favour stability in survival annual trends at southern latitudes (Yoccoz & Ims, 1999), as observed for *Apodemus* spp. in Italy, also when supplemental food was provided (P1.2b).

Conversely, trophic resources were the major driver of population abundance at both latitudes, with higher values in summer (P1.3a), and when supplemental food was provided (P1.3b). This might be linked to the restriction of home ranges when resources are abundant (see e.g. Stradiotto et al., 2009; Taitt & Krebs, 1981), which our models support by showing that individual residency time increased in proximity of food resources, when these were provided (Ferrari G., pers. comm.). As a consequence, the creation of vacant areas due to home range contraction might discourage emigration and favor immigration (Le Galliard et al., 2012; Rémy et al., 2013), ultimately increasing population size. Remarkable exceptions to this general pattern were recorded: in Norway, *M. glareolus* showed an overall decreasing trend in population size, due to the upcoming crash phase of multi-annual cycles (as detected in Johnsen et al., 2019) that typically govern rodents dynamics at northern latitudes (Andreassen et al., 2021); in Italy, population abundance of both *Apodemus* spp. and *M. glareolus* kept increasing from summer 2020 to winter 2020-2021, following a beech and spruce mast seeding, which produced overabundant spatially dispersed food resources (Bogdziewicz at al., 2016); further, population size of *M. glareolus* in Italian alpine woodlands exhibited an asynchronous seasonal pattern with respect to that of *Apodemus* spp., with peaks in winter and low phases in summer (P2.2a).

At southern latitudes, milder conditions in forestry environments promoted the co-occurrence of multiple species, in accordance with latidudinal biodiversity gradients (Araújo & Costa-Pereira, 2013; Mannion et al., 2014) (P2.1). While spring decline has been recorded in voles in other habitats (see Flowerdew et al., 1985 for a review), we intepret the asynchrony of annual population trends within the rodent community as an evidence of temporal niche partitioning. In sympatry, *Apodemus* spp. are generally dominant over *M. glareolus* (Amori et al., 2015; Casula et al., 2019), with the latter thus being forced to shift its niche to avoid competition. Benefiting by its wide dietary spectrum (Butet & Delettre, 2011), the generalist *M. glareolus* may exploit harsher seasonal conditions to temporally shift its niche, thus partitioning it with respect to the diet-specialist *Apodemus* spp. (Viviano et al., 2022), not able to cope with winter resource depletion. The overabundance of concentrated food of our experiment altered these processes, consistently enhancing the population demography of *Apodemus* spp., so buffering its population decline in winter (P1.3b). As a consequence, the ecological niches of *Apodemus* spp. and *M. glareolus* should have overlapped, and the niche partitioning failed. This eventually resulted in interspecific competition (P2.2b), with *M. glareolus* survival decreasing and population size collapsing. In nature, behavioral interference and exploitative competition (Wallace & Benbow, 2019) often co-exist, and their effects are difficult to separate (Schmidt et al., 2005; Shenbrot & Krasnov, 2002). Our experimental setting of food manipulation helps speculate the underpinning process to the observed demographic patterns. The presence of clumped food resources might have triggered behavioral interference, attracting both species (higher residency time, Ferrari G. pers. comm.), but favoring the dominant one. When food is confined, it becomes a defensible resource for species that cache food in larger hoards such as *Apodemus* spp. (Zwolak, Bogdziewicz, Wróbel, et al., 2016), potentially leading to aggressive interactions. These interactions might have chased *M. glareolus* away from the area surrounding the sites, eventually causing the observed collapse of population size (see e.g. Eccard & Ylönen, 2007). The concurrent decrease of survival experienced by *M. glareolus* might be instead imputed to exploitative competition (Gilad, 2008). As concentrated food resources enhance it abundance, *Apodemus* spp. might have consumed other resources than those provided at feeding sites, depriving *M. glareolus* also of other food sources. The lack of competition in presence of overabundant and spatially dispersed food resources i.e. masting, supports this explanation.

Under climate change. we expect more permissive environmental conditions for the rodent community in the boreal-temperate gradient, with a likely northward expansion of species’ distributional limits (Parmesan, 2006). The colonization of new habitats by opportunistic species e.g. *Apodemus* spp. would expand the spatial overlap with species such as *M. glareolus* which is well-adapted to cold environments (Butet and Delettre 2011, Markova et al. 2018). This scenario is supported by our latitudinal comparison, which suggests a tendency to niche partitioning as sympatry establishes (Bartolommei et al., 2018), or species competition to the disadvantage of adaptable but subordinate species.

Our assessment of the role of extrinsic and intrinsic factors, in interplay with biotic interactions, in shaping rodent demography, highlights the complexity of such relationships. Only by simultaneously controlling for several components of the ecological niche of the studied species (extrinsic top-down factors, i.e. climate severity, through latitudinal multi-population comparison; extrinsic bottom-up factors, i.e. resource availability, through food manipulation; intrinsic cycles, through seasonal variation; competition, through assessment of interactions between sympatric species) we have been able to evaluate their relative impact on the demography of each species. We suggest adopting an analogous approach when investigating other underpinning determinants of rodent demography patterns, such as parasitism and emergence of related diseases (Deter et al., 2008; Telfer et al., 2005), e.g. by comparing host-parasite-pathogens associations across environmental gradients or under different intensity of human disturbance.

## Acknowledgements

G. Ferrari was supported by Inland Norway of Applied Sciences Doctoral Program in Applied ecology and biotechnology and Fondazione Edmund Mach International Doctoral Program. F.C. contributed to this work partly under the IRD Fellowship 2021/2022 at Fondation IMéRA, Institute for Advanced Studies at Aix-Marseille Université. We are grateful to all the people involved in the rodent trapping and data collection, and especially Dal Farra S., Inama E. and Savazza S. We thank all the members of Applied Ecology and Animal Ecology Research Units and researchers from Evenstad Campus for their valuable suggestions.

## Authors’ contributions

G.F., O.D., V.T. and F.C. conceived the ideas and designed methodology; F.C., V.T. and O.D. equally supervised the work; G.F., V.T. and F.O. collected the data in Italy and K.J. collected the data in Norway; G.F. analysed the data with the support of O.D.. F.C., V.T.; G.F. led the writing of the manuscript, together with O.D., V.T., F.O., and F.C. All authors contributed critically to the drafts and gave final approval for publication.

## Data Availability Statement

Data are available from the Eurosmallmammals database (https://eurosmallmammals.fmach.it).

