## Supplemental Material for "Food resources drive rodent population demography mediated by seasonality and inter-specific competition"

### Supporting information

#### Appendix S1: Trapping sessions

In Norway the dataset included 26 primary occasions with two to seven secondary occasions from 2013 to 2015 (Table S1.1), while Italian trapping data consisted of 18 primary occasions with 3 secondary occasions each from 2019 to 2021 (see Table S1.2). Since the analysis used calendar months as temporal categorical variable, we decided to unify the secondary occasions of captures occurring in different sessions but within the same month (e.g. March 2019 in Italy and July 2013 in Norway).

**Table S1:** Primary and secondary capture occasions in Norway (**Table S1.1**) and Italy (**Table S1.2**). Grey-shaded occasions in Norway showed when food was provided at treatment grids (see main text).

| Season | Summer 2013 | | | | | Winter 2013/2014 | | | | |
|---|---|---|---|---|---|---|---|---|---|---|
| Primary occasion | 1 | | 2 | 3 | 4 | 5 | 6 | 7 | 8 | 9 |
| Secondary occ 1 | 03 Jul | 22 Jul | 10 Aug | 09 Sep | 09 Oct | 09 Nov | 09 Dec | 20 Jan | 17 Feb | 17 Mar |
| Secondary occ 2 | 04 Jul | 23 Jul | 14 Aug | 10 Sep | 10 Oct | 11 Nov | 10 Dec | 21 Jan | 18 Feb | 18 Mar |
| Secondary occ 3 | 05 Jul | 24 Jul | 15 Aug | | | 12 Nov | 11 Dec | | 19 Feb | 19 Mar |
| Secondary occ 4 | 10 Jul | | 16 Aug | | | 13 Nov | 12 Dec | | 20 Feb | 20 Mar |
| Secondary occ 5 | | | 17 Aug | | | 14 Nov | | | | |

| Season | Summer 2014 | | | | | | Winter 2014/2015 | | | | |
|---|---|---|---|---|---|---|---|---|---|---|---|
| Primary occasion | 10 | 11 | 12 | 13 | 14 | 16 | 17 | 18 | 19 | 20 | 21 |
| Secondary occ 1 | 22 Apr | 19 May | 23 Jun | 21 Jul | 18 Aug | 13 Oct | 03 Nov | 08 Dec | 12 Jan | 10 Feb | 18 Mar |
| Secondary occ 2 | 23 Apr | 20 May | 24 Jun | 22 Jul | 19 Aug | 14 Oct | 04 Nov | 09 Dec | 13 Jan | 11 Feb | 19 Mar |
| Secondary occ 3 | 24 Apr | 21 May | 25 Jun | 23 Jul | 20 Aug | 15 Oct | 05 Nov | 10 Dec | 14 Jan | 12 Feb | 20 Mar |
| Secondary occ 4 | 25 Apr | 22 May | 26 Jun | 24 Jul | 21 Aug | 16 Oct | 06 Nov | 11 Dec | 15 Jan | 13 Feb | |

| Season | Summer 2015 | | | | |
|---|---|---|---|---|---|
| Primary occasion | 22 | 23 | 24 | 25 | 26 |
| Secondary occ 1 | 14 Apr | 18 May | 16 Jun | 13 Jul | 11 Aug |
| Secondary occ 2 | 15 Apr | 19 May | 17 Jun | 14 Jul | 12 Aug |
| Secondary occ 3 | 16 Apr | 20 May | 18 Jun | 15 Jul | 13 Aug |
| Secondary occ 4 | | 21 May | 19 Jun | 16 Jul | 14 Aug |

**Table S1.1**

| Season | Winter 2018/2019 | | | Summer 2019 | |
|---|---|---|---|---|---|
| Primary occasion | 1 | 2 | | 3 | 4 |
| Secondary occ 1 | 19 Feb | 05 Mar | 19 Mar | 25 Jun | 27 Aug |
| Secondary occ 2 | 20 Feb | 06 Mar | 20 Mar | 26 Jun | 28 Aug |

| | | | | | | | | |
|---|---|---|---|---|---|---|---|---|
| Secondary occ 3 | | | 21 Feb | 07 Mar | 21 Mar | 27 Jun | 29 Aug | |
| | Winter 2019/2020 | | | | | Summer 2020 | | |
| Primary occasion | 5 | 6 | 7 | 8 | 9 | 10 | 11 | 12 |
| Secondary occ 1 | 12 Nov | 10 Dec | 14 Jan | 11 Feb | 04 Mar | 21 Apr | 30 Jun | 25 Aug |
| Secondary occ 2 | 13 Nov | 11 Dec | 15 Jan | 12 Feb | 05 Mar | 22 Apr | 01 Jul | 26 Aug |
| Secondary occ 3 | 14 Nov | 12 Dec | 16 Jan | 13 Feb | 06 Mar | 23 Apr | 02 Jul | 27 Aug |
| | Winter 2020/2021 | | | | | Summer 2021 | | |
| Primary occasion | 13 | 14 | 15 | 16 | 17 | 18 | | |
| Secondary occ 1 | 17 Nov | 15 Dec | 12 Jan | 23 Feb | 9 Mar | 7 Apr | | |
| Secondary occ 2 | 18 Nov | 16 Dec | 13 Jan | 24 Feb | 10 Mar | 8 Apr | | |
| Secondary occ 3 | 19 Nov | 17 Dec | 14 Jan | 25 Feb | 11 Mar | 9 Apr | | |

**Table S1.2**

### Appendix S2: Trapping grid design

Trapping grid design. The panel A shows the cross-shaped design with 16 traps used in Norway, while in the panel B the square design of 64 traps applied in Italy is represented.

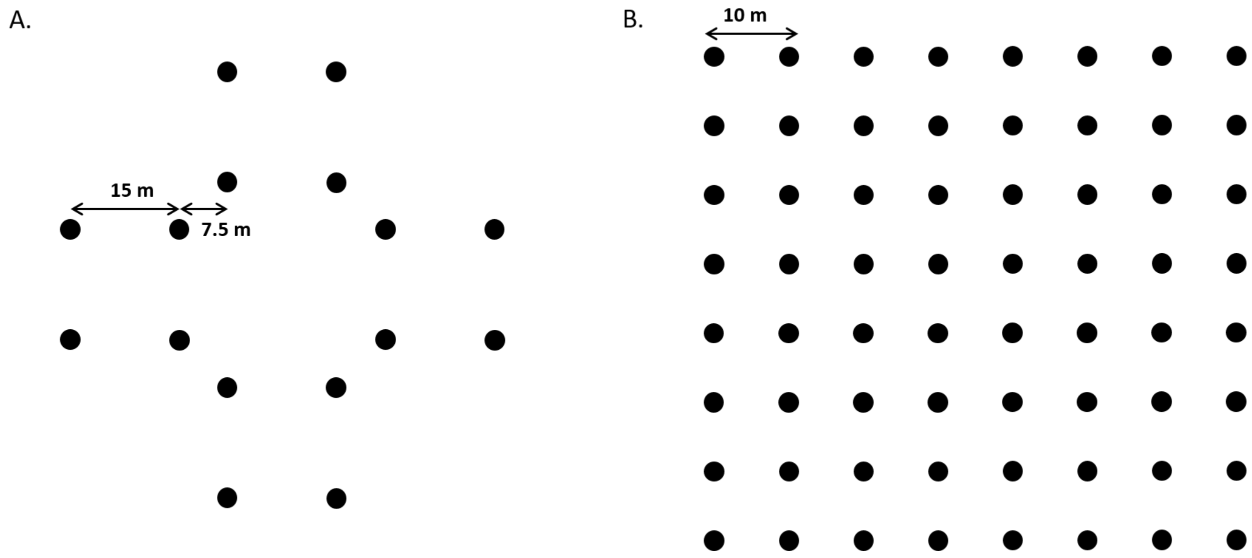

### Appendix S3: Preliminary exploratory analyses

#### Norway – Evenstad site

In Norway, as expected from the trapping protocol performed, only one species was encountered (*Myodes glareolus*). The density of captures had a similar pattern in females and males with two annual peaks during early summer and autumn (Figure S3.1; Table S3.1).

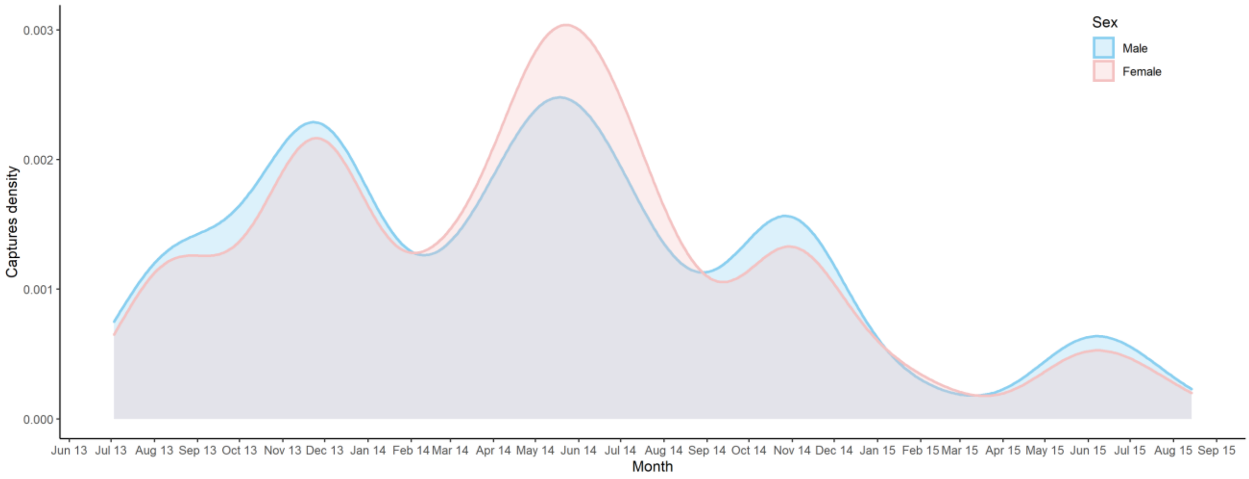


**Figure S3.1**: Captures density of *M. glareolus* in Norway across seasons in female (rose) and male (light blue).

#### Italy – Cembra site

When we compared the density of captures across trapping occasions, we detected a cyclic asynchronous pattern between the three detected species (*Apodemus* spp. and *Myodes glareolus*) (Figure S3.2). In particular, *Apodemus* spp. were captured more during summer compared to winter, with the exception of winter 2020-21. On the contrary, *M. glareolus* was captured more in winter than in summer.

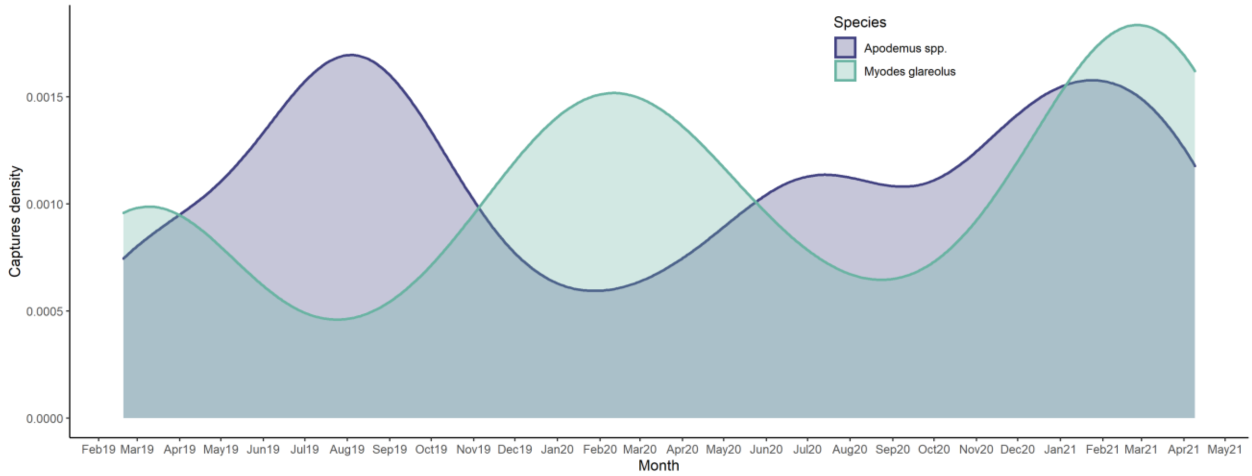


**Figure S3.2**: Capture density in Italy across seasons for *Apodemus* spp. (in purple) and *Myodes glareolus* (in green).

When breaking down the captures by sex, we found slightly shifted peaks for females and males in the three species, although the proportion of captures were similar (Figure S3.3; Table S3.1).

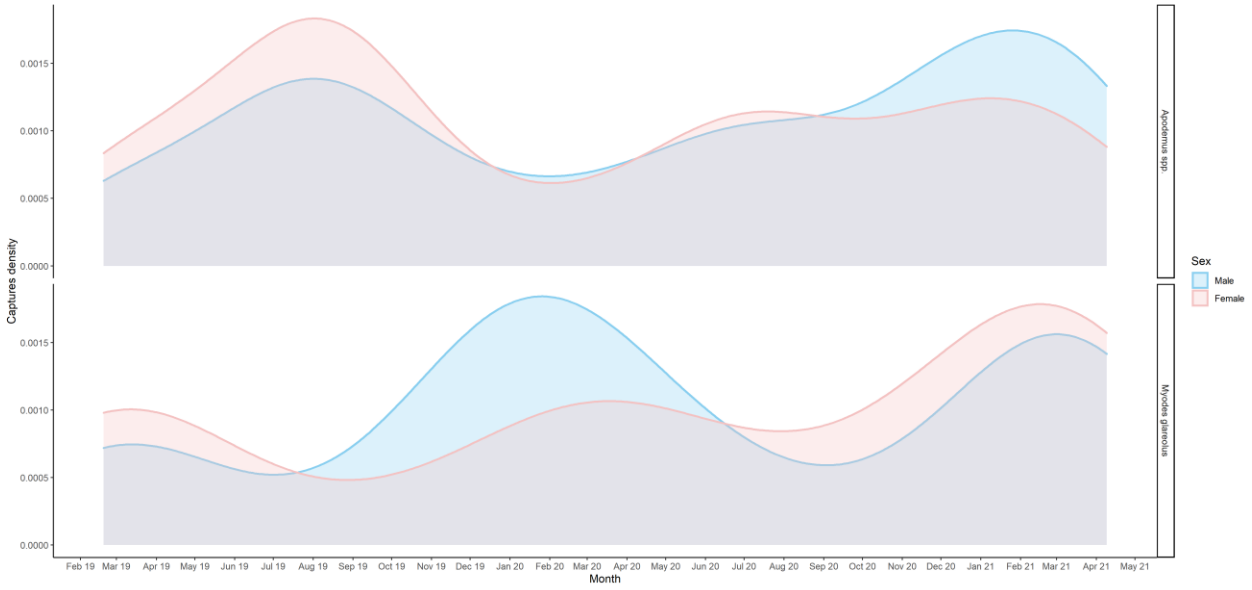


**Figure S3.3**: Captures density of *Apodemus* spp. and *Myodes glareolus* for female (rose) and male (light blue) individuals.

**Table S3.1**: Summary of the number of captures by sex in presence and absence of supplemental food for *M. glareolus* (Norway and Italy), and *Apodemus* spp. (Italy only).

| | *Myodes glareolus* | | *Apodemus spp.* | |
|---|---|---|---|---|
| Site | Feeding no | Feeding yes | Feeding no | Feeding yes |
| Norway | | | | |
| Individuals | 455 | 462 | | |
| Females | 212 | 213 | | |
| Males | 243 | 249 | | |
| Total captures | 1927 | 2049 | | |
| Italy | | | | |
| Individuals | 80 | 29 | 164 | 234 |
| Females | 34 | 10 | 84 | 117 |
| Males | 46 | 19 | 80 | 117 |
| Total captures | 391 | 62 | 440 | 486 |

#### Appendix S4: Models building

We identified a set of potentially biological meaningful covariates to disentangle seasonal, supplemental food availability, and inter-specific competition effect on rodent demography, and specifically:

(i) temporal covariates:

- **season (acting as an ordinary variable)**: successive binned time intervals based on seasonal periods throughout the study duration: winter (from November to March included) and summer (from April to October included);
- **time**: temporal effect within secondary occasions, i.e. daily **secondary trapping occasions;**
- **session**: temporal effect across primary occasions, i.e. monthly **primary trapping occasions**.

(ii) state/spatial covariate:

- **feeding state** of the site (Norway: spatio-temporally defined) and **feeding site** (Italy: spatially defined): presence (treatment) *vs* absence (control) of *ad libitum* supplemental feeding **(binary variables: yes/no)**.

(iii) individual covariate:

- **species (categorical variable;** Italy only**)**: *Apodemus* spp. and *Myodes glareolus*.

In CMR modelling, a main model is described by sub-models estimating each parameter, or response variables, in dependence on several covariates, or explanatory variables (Laake and Rexstad 2008). In particular, multistate open robust design (MSORD) is defined by five parameters (true survival rate ($S$), arrival probability (*pent*), apparent survival rate ($\varphi$), capture probability ($p$) and transition probability ($\psi$)).

For modelling each parameter, we chose those covariates that were biologically meaningful in our study systems, while being constrained by a relatively small sample size (i.e., few recaptures). In particular, we modelled true (*S*) and apparent survival (φ) as varying in dependence on primary occasions (‘session’, only for φ) and successive seasonal periods (‘season’) to detect the temporal pattern (H1), supplemental food availability (‘feeding state’ in Norway and ‘feeding site’ in Italy) to identify the food effect (H1), and species (only in Italy) to evaluate interspecific competition (H2). We considered the probabilities of capture (*p*) and arrival (*pent*) to be dependent on temporal variations (primary occasions i.e. ‘session’ for *p*, secondary occasions i.e. ‘time’ for *pent*), seasonal periods (‘season’), supplemental food availability (‘feeding state’-Norway and ‘feeding site’-Italy) and species (only in Italy). We modelled $\psi$ in dependence on transition of feeding state for Norway, and kept it constant for Italy (expressing the probability of transition of animal observability in the trapping grid). Table S4 summarizes the list of covariates or combination of covariates used to build the sets of sub-models (univariate, additive, or with two-way interactions among covariates) that were subsequently combined into main models.

Then we used a model selection based on AICc scores to rank the main models composed by sub-models and we retained the models with ΔAICc ≤ 4 as equally plausible (Burnham and Anderson 2002). Among these models, we chose the best model as the more biologically meaningful (Appendix S5).

**Table S4**: Covariates and combination of covariates used to build the set of sub-models for each demographic parameter in Norway (**Table S4.1**) and Italy (**Table S4.2**). Legend: ‘S’ = true survival; ‘pent’ = entry probability; ‘φ’ = apparent survival; ‘p’ = capture probability; ‘ψ’ = transition probability, set as constant in Italy and dependent on feeding states in Norway; ‘session’ = primary trapping occasions; ‘time’ = secondary trapping occasions; ‘feeding site’ = sites with supplemental food (only in Italy); ‘feeding state’ = supplemental food at sites (only in Norway); ‘feeding state : to feeding state’

= transition between feeding states (only in Norway); ‘species’ = rodent species (only for Italy); ‘season’ = seasonal variation across years.

| **Table S4.1** | S | pent | φ | p | ψ |
|---|---|---|---|---|---|
| feeding state : to feeding state | | | | | X |
| state : session | | | | | |
| feeding state : season | X | X | X | X | |
| feeding state + session | | X | X | X | |
| feeding state + season | X | X | X | X | |
| feeding state | X | X | X | X | |
| season | X | X | X | X | |
| session | X | X | X | X | |

| **Table S4.2** | S | pent | φ | p | ψ |
|---|---|---|---|---|---|
| feeding site: season : species | X | | | | |
| feeding site : season | | X | | X | |
| feeding site : season + species | X | | | | |
| feeding site : species + season | X | | | | |
| feeding site + season + species | X | | | | |
| feeding site + season | X | X | | X | |
| feeding site + session | | | | X | |
| feeding site + species | X | | | | |
| season + species | X | | | | |
| feeding site | X | X | X | X | |
| season | X | X | X | X | |
| session | X | | | X | |
| time | | X | | | |
| species | X | X | X | X | |
| 1 | X | X | X | X | X |

### Appendix S5: Model selection

In Norway, we found only one model with ΔAICc ≤ 4 (Model1). In the rest of the models, *S* and *pent* were dependent on ‘feeding state’, ‘session’ and ‘season’. Moreover, for *pent*, most of the models were defined by ‘feeding state : session’ and ‘session’ effect; while *φ* seemed equally affected by ‘feeding state’, and equally by ‘session’ and ‘season’. Finally, in mostly models, *p* was dependent on ‘feeding state’ and equally by ‘session’ and ‘season’ (Table S5.1). In conclusion, we decided to select Model1, because it showed the lowest AIC score and was the most biologically meaningful. Model1 retained ‘feeding state : session’ effect on *S*, ‘season + state’ on *pent* and finally ‘season’ on *φ* and ‘feeding state’ on *p*.

In Italy, the model selection returned 3 models with ΔAICc ≤ 4 which showed that ‘species’, ‘feeding site’, and ‘season had a stronger effect on *S*. In the three models, *pent* was always affected by ‘time’; *φ* by ‘species’ and *p* by ‘feeding site + session’ (Table S5.2). Although Model1 was the best parsimonious model with less parameters and lower AICc, we selected Model3 as best model because it allowed us to test our hypotheses. Specifically, Model 3 retained the interaction between ‘feeding site’ and ‘species’ added to ‘period’ on *S*; ‘time’ on *pent*; ‘species’ on *φ* and finally, ‘feeding + session’ on *p*.

**Table S5**: Model selection of Norwegian (**Table S5.1**) and Italian (**Table S5.2**) rodent population and capture parameters, reporting the best models (i.e. ΔAICc<4; in bold the one used for model predictions as most biologically meaningful with respect to the tested hypotheses), and the first model below the AICc threshold (in italic). Legend: 'S' = true survival; 'pent' = entry probability; 'φ' = apparent survival; 'p' = capture probability; 'ψ' = transition probability, set as constant in Italy and dependent on feeding states in Norway; 'session' = primary trapping occasions; 'time' = secondary trapping occasions; 'feeding site' = sites with supplemental food (only in Italy); 'feeding state' = supplemental food at sites (only in Norway); 'feeding state : to feeding state' = transition between feeding states (only in Norway); 'species' = rodent species (only for Italy); 'season' = seasonal variation across years; 'npar' = number of parameters, 'AICc' = AIC with a correction for small sample sizes; 'ΔAICc' = relative differences between the fitted model and the Akaike 'best-ranked' model with the smallest AICc value; 'AICc weight' = relative likelihood of a model; 'Deviance' = difference between null deviance and model deviance.

| **Table 5.1** | S | | | pent | | | Φ | | | p | | ψ | | | | | |
|---|---|---|---|---|---|---|---|---|---|---|---|---|---|---|---|---|---|
| Model Norway | feeding state | season | session | feeding state | season | session | season | session | feeding state | season | feeding state | feeding state : to feeding state | npar | AICc | ΔAICc | AICc weight | Deviance |
| **1** | **X** | **X** | | **X** | | **X** | **X** | | | | **X** | **X** | **45** | **8795.79** | **0.00** | **0.74** | **8704.61** |
| 2 | | | X | X | X | | | X | | X | X | X | 65 | 8797.86 | 2.07 | 0.26 | 8665.40 |
| *3* | *X* | *X* | | *X* | | *X* | | *X* | *X* | | *X* | *X* | *69* | *9631.72* | *835.93* | *0.00* | *9490.94* |

| **Table 5.2** | S | | | | pent | φ | | p | | ψ | | | | | |
|---|---|---|---|---|---|---|---|---|---|---|---|---|---|---|---|
| Model Italy | feeding site | species | season | feeding site : species | time | species | feeding site | feeding site | session | . | npar | AICc | ΔAICc | AICc weight | Deviance |
| 1 | | X | X | | X | X | | X | X | X | 33 | 11415.11 | 0 | 0.56 | 11347.41 |
| 2 | X | X | X | | X | X | | X | X | X | 34 | 11417.15 | 2.04 | 0.20 | 11347.35 |
| **3** | | | **X** | **X** | **X** | **X** | | **X** | **X** | **X** | **36** | **11417.69** | **2.58** | **0.15** | **11343.67** |
| *4* | | *X* | *X* | | *X* | | *X* | *X* | *X* | *X* | *33* | *11421.11* | *6.00* | *0.02* | *11353.42* |

### Appendix S6: Selected models' output: parameters' estimates

#### Norway – Evenstad site

**Table S6.1**: Parameter estimates of the best model (Model1, Table S5.1), with standard error (SE) and confidence intervals (Lower, Lcl, and Upper, Ucl). Legend: 'S' = true survival; 'pent' = entry probability; 'φ' = apparent survival; 'p' = capture probability; 'ψ' = transition probability depending on feeding states; 'session' = primary trapping occasions; 'feeding state' = supplemental food at sites; 'season' = seasonal variation across years. For some sessions, the model fitting did not converge.

| Parameter | Estimate | SE | Lcl | Ucl | Feeding state | Session | Season |
|---|---|---|---|---|---|---|---|
| S | 0.84 | 0.02 | 0.80 | 0.87 | NO | 1 | Summer2013 |
| S | 0.68 | 0.02 | 0.64 | 0.73 | NO | 5 | Winter2013-14 |
| S | 0.56 | 0.03 | 0.51 | 0.61 | NO | 10 | Summer2014 |
| S | 0.50 | 0.04 | 0.43 | 0.58 | NO | 17 | Winter2014-15 |
| S | 0.31 | 0.07 | 0.19 | 0.45 | NO | 22 | Summer2015 |
| S | 0.89 | 0.02 | 0.85 | 0.92 | YES | 1 | Summer2013 |
| S | 0.77 | 0.02 | 0.73 | 0.80 | YES | 5 | Winter2013-14 |
| S | 0.67 | 0.02 | 0.63 | 0.70 | YES | 10 | Summer2014 |
| S | 0.61 | 0.03 | 0.55 | 0.67 | YES | 17 | Winter2014-15 |
| S | 0.41 | 0.07 | 0.27 | 0.55 | YES | 22 | Summer2015 |
| S | NA | NA | NA | NA | Un-observable | 1 | Summer2013 |
| pent | 0.15 | 0.01 | 0.14 | 0.17 | NO | 1 | Summer2013 |
| pent | NA | NA | NA | NA | NO | 2 | Summer2013 |
| pent | 0.06 | 0.03 | 0.02 | 0.14 | NO | 3 | Summer2013 |
| pent | 0.56 | 0.06 | 0.44 | 0.67 | NO | 4 | Summer2013 |
| pent | 0.24 | 0.01 | 0.23 | 0.26 | NO | 5 | Winter2013-14 |
| pent | NA | NA | NA | NA | NO | 7 | Winter2013-14 |
| pent | NA | NA | NA | NA | NO | 16 | Summer2014 |
| pent | 0.17 | 0.16 | 0.02 | 0.64 | NO | 21 | Winter2014-15 |
| pent | NA | NA | NA | NA | NO | 22 | Summer2015 |
| pent | 0.17 | 0.10 | 0.05 | 0.44 | NO | 25 | Summer2015 |
| pent | NA | NA | NA | NA | YES | 3 | Summer2013 |
| pent | 0.23 | 0.11 | 0.08 | 0.51 | YES | 7 | Winter2013-14 |

| | | | | | | | |
|---|---|---|---|---|---|---|---|
| pent | NA | NA | NA | NA | YES | 8 | Winter2013-14 |
| pent | 0.33 | 0.00 | 0.33 | 0.33 | YES | 9 | Winter2013-14 |
| φ | 0.60 | 0.05 | 0.50 | 0.68 | NO | 1 | Summer2013 |
| φ | 0.78 | 0.02 | 0.73 | 0.83 | NO | 5 | Winter2013-14 |
| φ | 0.85 | 0.05 | 0.73 | 0.92 | NO | 10 | Summer2014 |
| φ | NA | NA | NA | NA | NO | 17 | Winter2014-15 |
| φ | 0.98 | 0.04 | 0.48 | 1.00 | NO | 22 | Summer2015 |
| p | 0.66 | 0.02 | 0.61 | 0.71 | NO | 1 | Summer2013 |
| p | 0.40 | 0.02 | 0.37 | 0.44 | YES | 1 | Summer2013 |
| ψ | 0.16 | 0.02 | 0.13 | 0.19 | NO; YES | 1 | Summer2013 |
| ψ | 0.03 | 0.02 | 0.01 | 0.10 | NO; Un-observable | 1 | Summer2013 |
| ψ | 0.06 | 0.01 | 0.05 | 0.08 | YES; NO | 1 | Summer2013 |
| ψ | NA | NA | NA | NA | YES; Un-observable | 1 | Summer2013 |
| ψ | NA | NA | NA | NA | Un-observable; NO | 1 | Summer2013 |
| ψ | NA | NA | NA | NA | Un-observable; YES | 1 | Summer2013 |

**Table S6.2**: Derived estimates of population size ($N_t$) retrieved by the best model (Model 1, Table S5.1), with confidence intervals (Lower, Lcl, and Upper, Ucl). Legend: ‘session’ = primary trapping occasions; ‘feeding state’ = supplemental food at sites. For some sessions, the model fitting did not converge.

| Estimate | Lcl | Ucl | Session | Feeding site |
|---|---|---|---|---|
| 98.96 | 94.16 | 103.76 | 1 | NO |
| 195.09 | 180.85 | 209.34 | 2 | NO |
| 119.56 | 111.01 | 128.11 | 3 | NO |
| 145.37 | 136.93 | 153.82 | 4 | NO |
| 172.97 | 166.84 | 179.10 | 5 | NO |
| 198.12 | 183.65 | 212.58 | 6 | NO |
| 68.06 | 63.09 | 73.03 | 13 | NO |
| 111.91 | 103.74 | 120.09 | 14 | NO |
| 108.67 | 104.72 | 112.63 | 15 | NO |
| 48.40 | 44.86 | 51.93 | 17 | NO |
| 19.66 | 18.23 | 21.10 | 18 | NO |
| 16.64 | 15.42 | 17.85 | 19 | NO |

| | | | | |
|---|---|---|---|---|
| 2.67 | 2.08 | 3.26 | 20 | NO |
| 3.27 | 3.15 | 3.39 | 21 | NO |
| 4.54 | 4.21 | 4.87 | 23 | NO |
| 4.98 | 3.98 | 5.98 | 24 | NO |
| 161.82 | 148.45 | 175.19 | 6 | YES |
| 141.76 | 131.05 | 152.48 | 7 | YES |
| 143.33 | 135.52 | 151.14 | 9 | YES |
| 497.91 | 456.78 | 539.04 | 10 | YES |
| 433.18 | 397.40 | 468.97 | 11 | YES |
| 351.03 | 322.03 | 380.02 | 12 | YES |
| 303.72 | 278.63 | 328.82 | 16 | YES |

#### Italy – Cembra site

**Table S6.3**: Parameter estimates of the best model (Model3, Table S5.2), with standard error (SE) and confidence intervals (Lower, Lcl, and Upper, Ucl). Legend: ‘S’ = true survival; ‘pent’ = entry probability; ‘φ’ = apparent survival; ‘p’ = capture probability; ‘ψ’ = transition probability set as constant; ‘session’ = primary trapping occasions; ‘feeding site’ = sites with supplemental food; ‘species’ = rodent species; ‘season’ = seasonal variation across years. For some sessions, the model fitting did not converge.

| Parameter | Estimate | SE | Lcl | Ucl | Session | Feeding site | Species | Season |
|---|---|---|---|---|---|---|---|---|
| S | 0.66 | 0.06 | 0.53 | 0.77 | 0 | NO | Apodemus spp. | Winter2018-19 |
| S | 0.44 | 0.05 | 0.36 | 0.54 | 4 | NO | Apodemus spp. | Summer2019 |
| S | 0.65 | 0.05 | 0.54 | 0.75 | 9 | NO | Apodemus spp. | Summer2019 |
| S | 0.52 | 0.05 | 0.43 | 0.61 | 14 | NO | Apodemus spp. | Winter2019-20 |
| S | 0.73 | 0.04 | 0.65 | 0.79 | 21 | NO | Apodemus spp. | Summer2020 |
| S | 0.69 | 0.06 | 0.57 | 0.80 | 0 | YES | Apodemus spp. | Winter2018-19 |
| S | 0.48 | 0.04 | 0.40 | 0.56 | 4 | YES | Apodemus spp. | Summer2019 |
| S | 0.69 | 0.05 | 0.59 | 0.77 | 9 | YES | Apodemus spp. | Summer2019 |
| S | 0.56 | 0.04 | 0.47 | 0.64 | 14 | YES | Apodemus spp. | Winter2019-20 |
| S | 0.76 | 0.03 | 0.68 | 0.82 | 21 | YES | Apodemus spp. | Summer2020 |
| S | 0.80 | 0.05 | 0.69 | 0.88 | 0 | NO | Myodes glareolus | Winter2018-19 |
| S | 0.63 | 0.05 | 0.52 | 0.72 | 4 | NO | Myodes glareolus | Summer2019 |
| S | 0.80 | 0.04 | 0.72 | 0.86 | 9 | NO | Myodes glareolus | Summer2019 |
| S | 0.69 | 0.04 | 0.60 | 0.77 | 14 | NO | Myodes glareolus | Winter2019-20 |
| S | 0.85 | 0.03 | 0.78 | 0.90 | 21 | NO | Myodes glareolus | Summer2020 |
| S | 0.63 | 0.11 | 0.41 | 0.82 | 0 | YES | Myodes glareolus | Winter2018-19 |
| S | 0.42 | 0.11 | 0.22 | 0.64 | 4 | YES | Myodes glareolus | Summer2019 |
| S | 0.63 | 0.11 | 0.40 | 0.81 | 9 | YES | Myodes glareolus | Summer2019 |
| S | 0.49 | 0.12 | 0.28 | 0.71 | 14 | YES | Myodes glareolus | Winter2019-20 |
| S | 0.71 | 0.09 | 0.50 | 0.85 | 21 | YES | Myodes glareolus | Summer2020 |
| pent | 0.78 | 0.04 | 0.70 | 0.84 | 1 | NO | Apodemus spp. | Winter2018-19 |
| pent | 0.10 | 0.04 | 0.05 | 0.20 | 1 | NO | Apodemus spp. | Winter2018-19 |
| pent | 0.61 | 0.07 | 0.47 | 0.73 | 2 | NO | Apodemus spp. | Winter2018-19 |
| pent | 0.08 | 0.03 | 0.04 | 0.16 | 2 | NO | Apodemus spp. | Winter2018-19 |

| | | | | | | | | |
|---|---|---|---|---|---|---|---|---|
| pent | 0.10 | 0.07 | 0.02 | 0.35 | 2 | NO | Apodemus spp. | Winter2018-19 |
| pent | 0.10 | 0.07 | 0.02 | 0.34 | 2 | NO | Apodemus spp. | Winter2018-19 |
| pent | 0.02 | 0.04 | 0.00 | 0.65 | 2 | NO | Apodemus spp. | Winter2018-19 |
| φ | 0.87 | 0.03 | 0.81 | 0.92 | 1 | NO | Apodemus spp. | Winter2018-19 |
| φ | 0.98 | 0.02 | 0.86 | 1.00 | 1 | NO | Myodes glareolus | Winter2018-19 |
| p | 0.77 | 0.06 | 0.63 | 0.87 | 1 | NO | Apodemus spp. | Winter2018-19 |
| p | 0.78 | 0.05 | 0.68 | 0.86 | 2 | NO | Apodemus spp. | Winter2018-19 |
| p | 0.56 | 0.06 | 0.43 | 0.67 | 5 | NO | Apodemus spp. | Summer2019 |
| p | 0.71 | 0.05 | 0.61 | 0.79 | 7 | NO | Apodemus spp. | Summer2019 |
| p | 0.56 | 0.07 | 0.41 | 0.69 | 10 | NO | Apodemus spp. | Winter2019-20 |
| p | 0.51 | 0.07 | 0.38 | 0.64 | 11 | NO | Apodemus spp. | Winter2019-20 |
| p | 0.39 | 0.07 | 0.26 | 0.53 | 12 | NO | Apodemus spp. | Winter2019-20 |
| p | 0.56 | 0.07 | 0.43 | 0.68 | 13 | NO | Apodemus spp. | Winter2019-20 |
| p | 0.61 | 0.07 | 0.46 | 0.73 | 14 | NO | Apodemus spp. | Winter2019-20 |
| p | 0.73 | 0.06 | 0.59 | 0.83 | 15 | NO | Apodemus spp. | Summer2020 |
| p | 0.78 | 0.04 | 0.68 | 0.86 | 17 | NO | Apodemus spp. | Summer2020 |
| p | 0.71 | 0.07 | 0.56 | 0.83 | 19 | NO | Apodemus spp. | Summer2020 |
| p | 0.50 | 0.07 | 0.37 | 0.63 | 22 | NO | Apodemus spp. | Winter2020-21 |
| p | 0.65 | 0.05 | 0.54 | 0.74 | 23 | NO | Apodemus spp. | Winter2020-21 |
| p | 0.18 | 0.04 | 0.12 | 0.28 | 24 | NO | Apodemus spp. | Winter2020-21 |
| p | 0.53 | 0.05 | 0.42 | 0.63 | 25 | NO | Apodemus spp. | Winter2020-21 |
| p | 0.60 | 0.05 | 0.50 | 0.69 | 26 | NO | Apodemus spp. | Winter2020-21 |
| p | 0.59 | 0.05 | 0.48 | 0.69 | 27 | NO | Apodemus spp. | Summer2021 |
| p | 0.56 | 0.09 | 0.39 | 0.72 | 1 | YES | Apodemus spp. | Winter2018-19 |
| p | 0.57 | 0.07 | 0.43 | 0.70 | 2 | YES | Apodemus spp. | Winter2018-19 |
| p | 0.32 | 0.05 | 0.22 | 0.43 | 5 | YES | Apodemus spp. | Summer2019 |
| p | 0.48 | 0.06 | 0.37 | 0.59 | 7 | YES | Apodemus spp. | Summer2019 |
| p | 0.32 | 0.06 | 0.21 | 0.45 | 10 | YES | Apodemus spp. | Winter2019-20 |
| p | 0.28 | 0.06 | 0.18 | 0.41 | 11 | YES | Apodemus spp. | Winter2019-20 |
| p | 0.19 | 0.05 | 0.11 | 0.30 | 12 | YES | Apodemus spp. | Winter2019-20 |
| p | 0.32 | 0.06 | 0.21 | 0.45 | 13 | YES | Apodemus spp. | Winter2019-20 |
| p | 0.36 | 0.07 | 0.24 | 0.50 | 14 | YES | Apodemus spp. | Winter2019-20 |
| p | 0.49 | 0.08 | 0.34 | 0.65 | 15 | YES | Apodemus spp. | Summer2020 |
| p | 0.57 | 0.06 | 0.45 | 0.68 | 17 | YES | Apodemus spp. | Summer2020 |
| p | 0.48 | 0.08 | 0.32 | 0.64 | 19 | YES | Apodemus spp. | Summer2020 |

| | | | | | | | | |
|---|---|---|---|---|---|---|---|---|
| p | 0.27 | 0.06 | 0.17 | 0.39 | 22 | YES | Apodemus spp. | Winter2020-21 |
| p | 0.40 | 0.06 | 0.30 | 0.52 | 23 | YES | Apodemus spp. | Winter2020-21 |
| p | 0.08 | 0.02 | 0.05 | 0.13 | 24 | YES | Apodemus spp. | Winter2020-21 |
| p | 0.29 | 0.05 | 0.21 | 0.39 | 25 | YES | Apodemus spp. | Winter2020-21 |
| p | 0.36 | 0.05 | 0.27 | 0.46 | 26 | YES | Apodemus spp. | Winter2020-21 |
| p | 0.35 | 0.05 | 0.26 | 0.45 | 27 | YES | Apodemus spp. | Summer2021 |
| ψ | NA | NA | NA | NA | 1 | NO | Apodemus spp. | Winter2018-19 |

**Table S6.4**: Derived estimates of population size ($N_t$) retrieved by the best model (Model3, Table S5.2) with confidence intervals (Lower, Lcl, and Upper, Ucl). Legend: 'S' = true survival; 'pent' = entry probability; 'φ' = apparent survival; 'p' = capture probability; 'ψ' = transition probability set as constant; 'session' = primary trapping occasions; 'feeding site' = sites with supplemental food; 'species' = rodent species. For some sessions, the model fitting did not converge.

| Estimate | Lcl | Ucl | Session | Feeding site | Species |
|---|---|---|---|---|---|
| 7.51 | 7.12 | 7.89 | 1 | NO | Apodemus spp. |
| 13.71 | 13.32 | 14.11 | 2 | NO | Apodemus spp. |
| 34.52 | 30.84 | 38.20 | 3 | NO | Apodemus spp. |
| 34.25 | 32.64 | 35.85 | 4 | NO | Apodemus spp. |
| 3.69 | 3.21 | 4.17 | 5 | NO | Apodemus spp. |
| 2.57 | 2.20 | 2.94 | 6 | NO | Apodemus spp. |
| 1.52 | 1.17 | 1.87 | 7 | NO | Apodemus spp. |
| 4.91 | 4.33 | 5.49 | 8 | NO | Apodemus spp. |
| 5.91 | 5.30 | 6.52 | 9 | NO | Apodemus spp. |
| 8.78 | 8.24 | 9.31 | 10 | NO | Apodemus spp. |
| 21.38 | 20.61 | 22.16 | 11 | NO | Apodemus spp. |
| 13.26 | 12.34 | 14.19 | 12 | NO | Apodemus spp. |
| 27.43 | 23.36 | 31.51 | 13 | NO | Apodemus spp. |
| 21.82 | 20.34 | 23.31 | 14 | NO | Apodemus spp. |
| 18.71 | 11.94 | 25.47 | 15 | NO | Apodemus spp. |
| 24.11 | 21.56 | 26.66 | 16 | NO | Apodemus spp. |
| 14.25 | 13.18 | 15.33 | 17 | NO | Apodemus spp. |
| 27.57 | 25.31 | 29.82 | 18 | NO | Apodemus spp. |

| | | | | | |
|---|---|---|---|---|---|
| NA | NA | NA | 1 | NO | Apodemus spp. |
| 6.14 | 5.21 | 7.07 | 1 | YES | Apodemus spp. |
| 13.77 | 12.75 | 14.79 | 2 | YES | Apodemus spp. |
| 84.05 | 65.12 | 102.99 | 3 | YES | Apodemus spp. |
| 61.51 | 53.79 | 69.24 | 4 | YES | Apodemus spp. |
| 22.69 | 16.27 | 29.11 | 5 | YES | Apodemus spp. |
| 15.31 | 10.66 | 19.96 | 6 | YES | Apodemus spp. |
| 7.82 | 4.67 | 10.96 | 7 | YES | Apodemus spp. |
| 5.21 | 3.78 | 6.63 | 8 | YES | Apodemus spp. |
| 7.95 | 5.96 | 9.94 | 9 | YES | Apodemus spp. |
| 19.68 | 16.29 | 23.06 | 10 | YES | Apodemus spp. |
| 41.40 | 37.20 | 45.60 | 11 | YES | Apodemus spp. |
| 18.75 | 15.14 | 22.35 | 12 | YES | Apodemus spp. |
| 43.48 | 30.21 | 56.76 | 13 | YES | Apodemus spp. |
| 26.69 | 22.04 | 31.33 | 14 | YES | Apodemus spp. |
| 46.52 | 24.34 | 68.70 | 15 | YES | Apodemus spp. |
| 14.90 | 11.44 | 18.36 | 16 | YES | Apodemus spp. |
| 32.17 | 26.46 | 37.88 | 17 | YES | Apodemus spp. |
| 34.50 | 28.09 | 40.91 | 18 | YES | Apodemus spp. |
| 8.38 | 8.05 | 8.71 | 1 | NO | Myodes glareolus |
| 10.29 | 10.11 | 10.46 | 2 | NO | Myodes glareolus |
| 7.06 | 6.38 | 7.74 | 3 | NO | Myodes glareolus |
| 7.51 | 7.22 | 7.79 | 4 | NO | Myodes glareolus |
| 5.88 | 5.20 | 6.55 | 5 | NO | Myodes glareolus |
| 14.66 | 12.80 | 16.53 | 6 | NO | Myodes glareolus |
| 12.84 | 10.08 | 15.59 | 7 | NO | Myodes glareolus |
| 15.25 | 13.66 | 16.83 | 8 | NO | Myodes glareolus |
| 10.21 | 9.29 | 11.12 | 9 | NO | Myodes glareolus |
| 13.87 | 13.19 | 14.54 | 10 | NO | Myodes glareolus |
| 8.37 | 8.13 | 8.60 | 11 | NO | Myodes glareolus |
| 6.44 | 6.07 | 6.80 | 12 | NO | Myodes glareolus |
| 8.68 | 7.50 | 9.85 | 13 | NO | Myodes glareolus |
| 11.07 | 10.43 | 11.72 | 14 | NO | Myodes glareolus |
| 24.63 | 15.87 | 33.39 | 15 | NO | Myodes glareolus |
| 14.49 | 13.09 | 15.89 | 16 | NO | Myodes glareolus |

| | | | | | |
|---|---|---|---|---|---|
| 19.36 | 18.10 | 20.63 | 17 | NO | Myodes glareolus |
| 16.07 | 14.92 | 17.23 | 18 | NO | Myodes glareolus |
| 5.86 | 5.07 | 6.65 | 1 | YES | Myodes glareolus |
| 3.23 | 3.08 | 3.39 | 2 | YES | Myodes glareolus |
| 1.63 | 1.18 | 2.07 | 5 | YES | Myodes glareolus |
| 2.40 | 1.45 | 3.35 | 7 | YES | Myodes glareolus |
| 2.98 | 2.27 | 3.69 | 9 | YES | Myodes glareolus |
| 1.24 | 1.05 | 1.44 | 10 | YES | Myodes glareolus |
| 1.39 | 1.16 | 1.63 | 14 | YES | Myodes glareolus |
| 5.20 | 4.01 | 6.39 | 16 | YES | Myodes glareolus |
| 16.56 | 13.73 | 19.38 | 17 | YES | Myodes glareolus |
| 21.50 | 17.63 | 25.37 | 18 | YES | Myodes glareolus |

#### Appendix S7: Ancillary demographic parameters

*Apparent survival*

Apparent survival ($\varphi$) showed differences across the two latitudes. In particular, in Norway $\varphi$ in *M. glareolus* varied through seasons with slight differences between winter and summer, but no feeding effect was detected (Figure S7.1).

In Italy, on the contrary, $\varphi$ was affected by species and it was higher in *M. glareolus* than in *Apodemus spp.* (Figure S7.2).

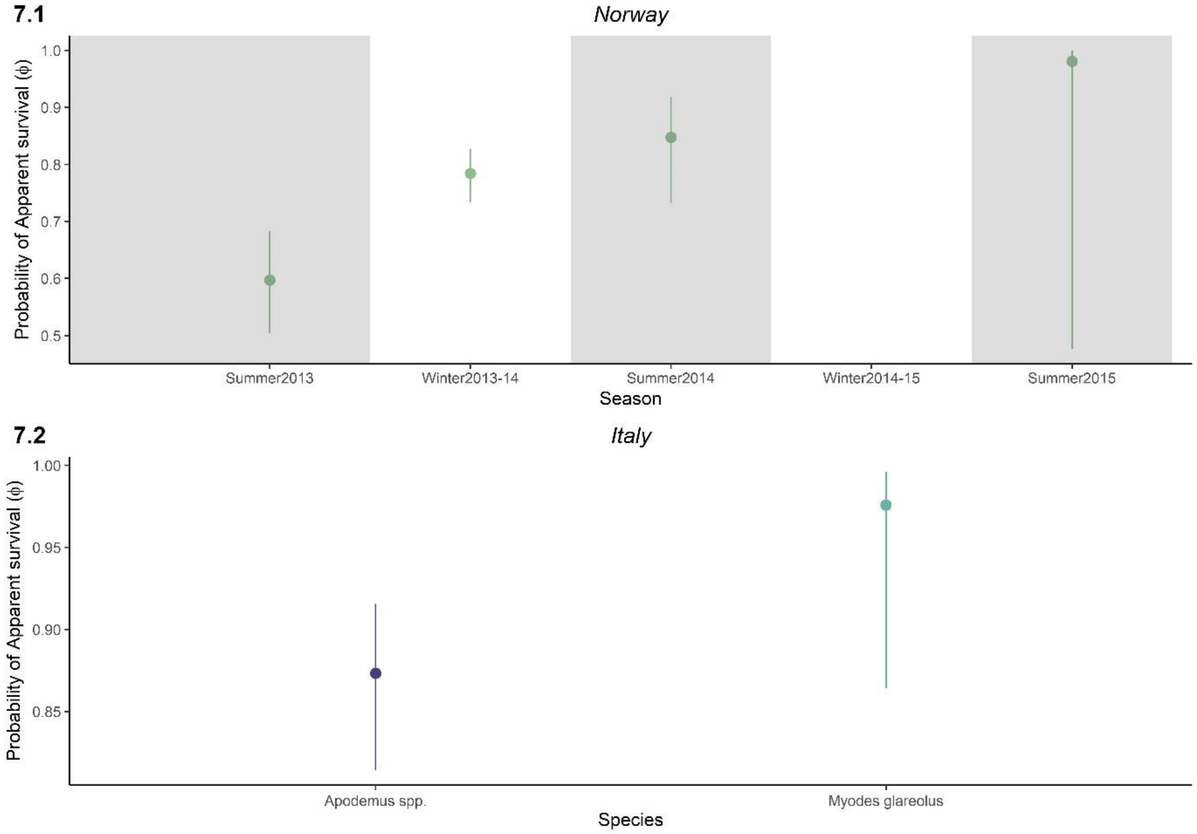


**Figure S7.1 and S7.2**: Real estimates of apparent survival (φ) in Norway (**Figure S7.1**, above) depending on seasonal periods and in Italy (**Figure S7.2**, below) for each detected species (*M. glareolus* and *Apodemus spp.*). Grey-shaded boxes show the summer periods, while white boxes show the winter periods.

*Arrival probability*

Arrival probability (*pent*) was time-dependent both in Norway and in Italy. In Norway *pent* retained a temporal effect, but depended on primary occasions rather than on secondary occasions (Figure S7.3). In particular, *pent* was high during autumn when food was not provided, and then during winter with supplemental food.

In Italy *pent* was affected by secondary trapping occasions, with higher values at the first secondary occasion, which decreased with the subsequent fives (Figure S7.4).

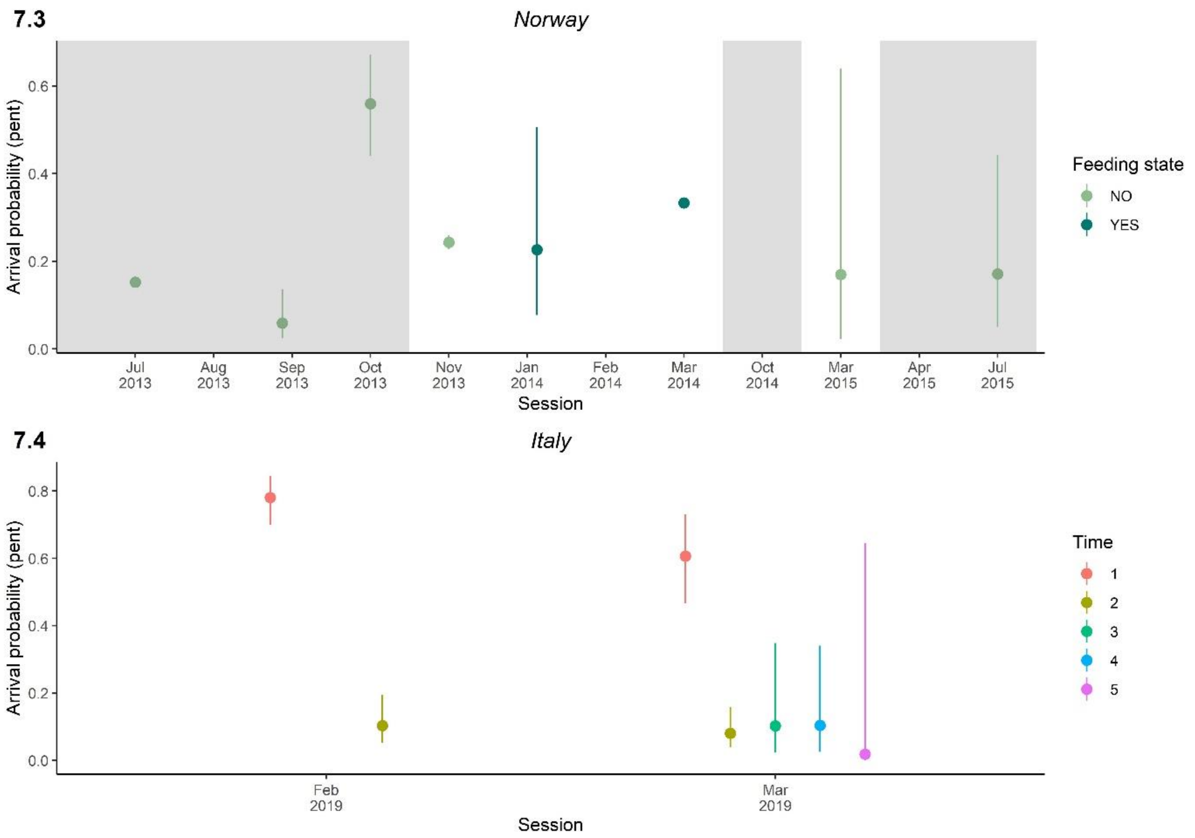


**Figure S7.3 and S7.4**: Real estimates of arrival probability (pent) in Norway (**Figure S7.3**) for each seasonal periods and under treatment (supplemental food availability; dark green) and control (no supplemental food; light green) conditions, and in Italy (**Figure S7.4**) for each secondary trapping occasions. Grey-shaded boxes show the summer periods, while white boxes show the winter periods.

*Capture probability*

In Norway, *p* depended only on feeding state and in particular, when supplemental food was provided, animals were less captured than without food (Figure S7.5).

Similarly, in Italy *p* changed in relation with supplemental food and across primary occasions. In particular, *p* was lower where food was provided during the entire sampling period for both species. In general, *p* declined from summer to winter, with some exceptions (December 2020 and January 2021) (Figure S7.6).

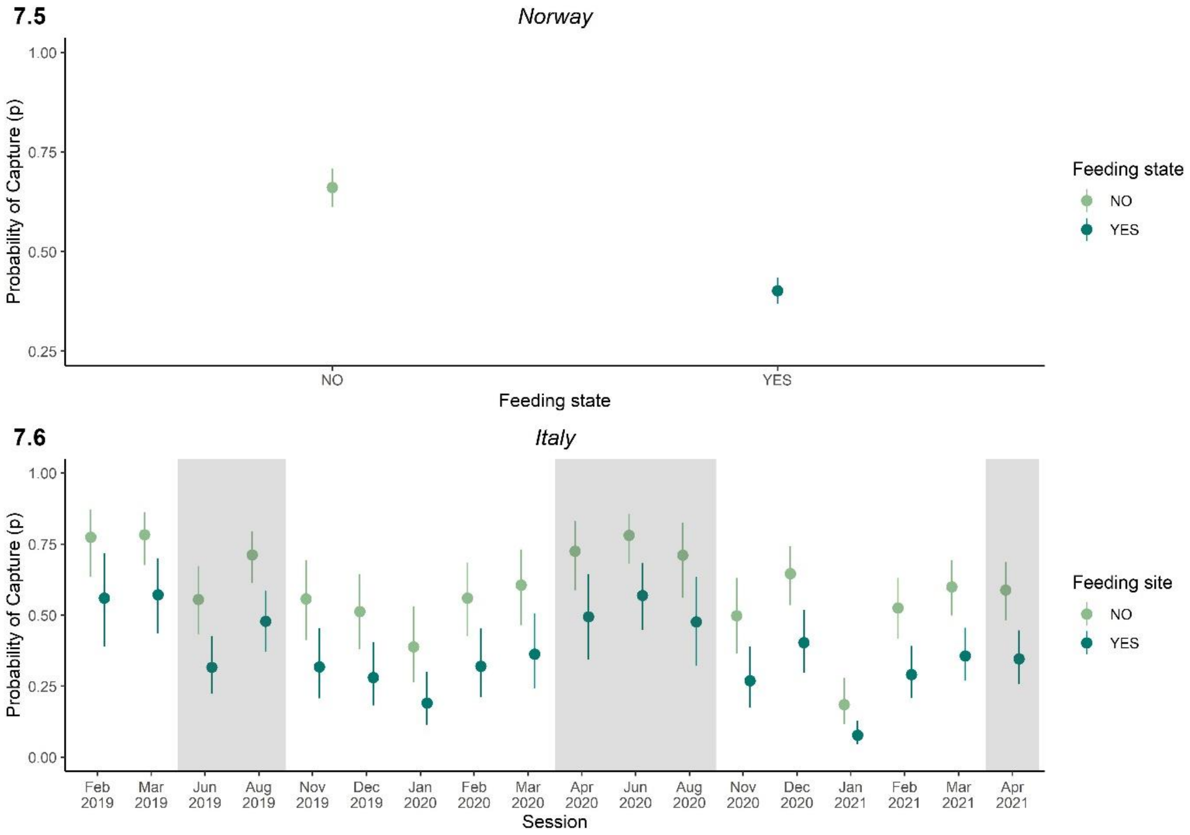


**Figure S7.5 and S7.6**: Estimates of capture probability (*p*) in rodents in Norway (**Figure S7.5**) and in Italy (**Figure S7.6**) for each primary trapping occasion, with treatment supplemental food (supplemental food available; dark green) and control (no supplemental food available; light green) conditions (‘Feeding state’ for Norway and ‘Feeding site’ for Italy). Grey-shaded boxes show the summer periods, while white boxes show the winter periods.
